# Stable neuronal representations to repeated stimulation underlie cognitive resilience in Alzheimer’s disease pathology

**DOI:** 10.64898/2025.12.05.692431

**Authors:** Keying Chen, Emma Pineau, Margaret Koletar, Andrea Trevisiol, Jack Simiao He, Mary Hill, Maged Goubran, John Sled, JoAnne McLaurin, Bojana Stefanovic

**Affiliations:** Physical Sciences, Sunnybrook Research Institute, Toronto, ON, Canada; Hurvitz Brain Sciences, Sunnybrook Health Sciences Centre, Toronto, Ontario, Canada; Department of Medical Biophysics, University of Toronto, Toronto, ON, Canada; Faculty of Health Sciences, McMaster University; Biological Sciences, Sunnybrook Research Institute, Toronto, ON, Canada; Department of Laboratory Medicine and Pathobiology, Temerty Faculty of Medicine, University of Toronto, Toronto, ON, Canada; Harquail Centre for Neuromodulation, Sunnybrook Health Sciences Centre, Toronto, Ontario, Canada; Mouse Imaging Centre, The Hospital for Sick Children, Toronto, ON, Canada; Translational Medicine, SickKids Research Institute, Toronto, ON, Canada; Department of Obstetrics & Gynecology, University of Toronto, Toronto, ON, Canada

## Abstract

While Alzheimer’s Disease (AD) typically triggers cognitive decline, some individuals with significant AD pathology maintain normal cognition into late life. Understanding the neuronal underpinnings of such cognitive resilience would propel the development of interventions for delaying dementia. To this end, we used cognitive testing to identify a subset of cognitively resilient 13-month-old TgF344-AD rats (established AD) and their non-transgenic littermates, followed by Neuropixels recording from 8500 neurons during repeated somatosensory stimulation. Cognitively resilient TgF344-AD rats recruited fewer neurons yet displayed more stable neuronal representations during repeated stimulations in cortical excitatory and hippocampal inhibitory ensembles, with reduced excitatory spike burstiness during network activation and a distinct pattern of functional synaptic connectivity. These associations existed independently of amyloid and tau levels. For the first time, our study revealed neuronal population-level hallmarks of maintained cognition that may serve as a novel neurophysiological biomarker of cognitive resilience and a target for stabilizing cognition.

## Introduction

Alzheimer’s disease (AD) as the most common cause of dementia is pathologically characterized by amyloid-β plaques and tau neurofibrillary tangles^1,2^. However, cognitive impairment may not necessarily result from either amyloid-β or tau accumulation ^3–6^, as up to one-third of elderly maintain intact cognitive performance despite substantial AD pathology ^7,8^. This phenomenon of cognitive resilience in AD has attracted much research interest as it may provide clues for delaying or preventing dementia ^9^. Current knowledge of resilience largely stems from large-scale measures such as cortical thickness, functional connectivity and structural network organization ^10–20^, as well as single-cell and multi-omics studies that probe the molecular signatures of resilience ^21–24^. In contrast, circuit-level mechanisms remain poorly understood; to date, only Shoob et al. have identified circuit-level homeostasis in APP/PS1 mice as one correlate of resilience ^25^. To identify candidate neurophysiological biomarkers and neuromodulatory intervention targets to preserve cognitive function, there is an urgent need for *in vivo*, high-spatiotemporal-resolution mapping of the functional networks that support cognitive resilience in the context of AD pathology.

A prominent theory regarding cognitive reserve posits that the onset of dementia depends on both passive reserve (the quantity of neurons and synapses) and active reserve (the brain’s capacity to reorganize processing, e.g. through recruitment of alternative circuits) ^26,27^. Thus, it is critical to reveal differences in how resilient vs. non-resilient AD brains allocate neuronal resources, and how these patterns differ from those observed in healthy aging. In healthy brains, neuronal responses to repeated stimulation are highly variable, with only 15–25% overlap in the subset of hippocampal neurons activated across trials ^28,29^. This variability indicates that the same stimulus can be processed by multiple, only partially overlapping neuronal ensembles. Interestingly, computational modeling studies indicate that although individual neurons may be highly sensitive to certain stimuli, most of the information processing is carried out by overlapping neuronal populations ^30–32^. These computationally-derived conclusions are supported by experimental studies in healthy brains ^33,34^. However, it remains unknown whether the AD brain can reliably recruit a stable neuronal population across repeated stimulation, and how this reliability may be linked to resilience.

Advances in neurotechnology enabled tracking hundreds of neurons simultaneously with millisecond precision across brain regions using state-of-the-art ultra-high-density electrophysiology probes. The novelty of our study lies in leveraging this technique to connect single-neuron activity with microcircuit organization in the context of cognitive resilience in AD. We here defined cognitive resilience as behavioral performance of 13-month-old rats comparable to that of a young 4-month-old reference cohort. While age-matched non-transgenic littermates (nTg) served as controls for non-diseased aging, the AD transgenic Fischer-344 rats (TgF344-AD; “TgAD”) is well suited for resilience studies because it shows substantial amyloid burden, tau pathology, and cognitive decline corresponding to early symptomatic Alzheimer’s disease, and offers greater translational relevance than most transgenic AD mouse models ^35,36^. We provide the first *in vivo*, cellular-resolution characterization of population-level circuit dynamics underlying resilience in a rat model of AD. We found that cognitively unimpaired TgAD rats (“AD Resilient”), similar to cognitively unimpaired nTg rats (“Healthy Aged”), exhibited more stable population-level recruitment of cortical excitatory neurons and hippocampal inhibitory neurons during repeated forepaw stimulation than did cognitively impaired TgAD (“AD Impaired”) and nTg rats (“Aging Impaired”). Despite the presence of AD pathologies, “AD Resilient” Tg rats preserved network function by preventing excessive hyperexcitability through inhibitory-to-inhibitory synaptic connection number, strength, and divergence. These findings provide a cellular-scale understanding of neuronal networks function and connectivity that underlie cognitive resilience in AD pathologies. This work identifies neurophysiological biomarkers to use in the development of interventions for preserving brain function in the face of progressive neurodegeneration.

## Results

In this study (**Fig.1**), we identified a subset of 13-month-old TgAD rats that maintained cognitive performance despite exhibiting amyloid burden that was comparable to that of their cognitively impaired littermates. Electrophysiological recordings from these rats during repeated transcutaneous forepaw stimulation allowed us to examine how neurons coordinate their activity in the context of cognitive resilience in Alzheimer’s disease (AD) pathologies and/or aging. We organized the electrophysiological results across spatial scales. We first quantified population-level neural activity by calculating local field potential (LFP) power. Then we focused on single-unit (SU) recordings to capture individual neurons’ activity. After assessing the SU signal quality, we analyzed neuronal responses to sensory forepaw stimulation, examining the number of neurons activating, firing rates, inter-spike interval patterns, and neuronal engagement at the network level. Subsequently, we analyzed synaptic connectivity to investigate whether distinct synaptic patterns explain the observed activity differences. Together, our results yielded a comprehensive understanding of cognitive resilience at the early symptomatic stage of AD.

**Figure 1:**
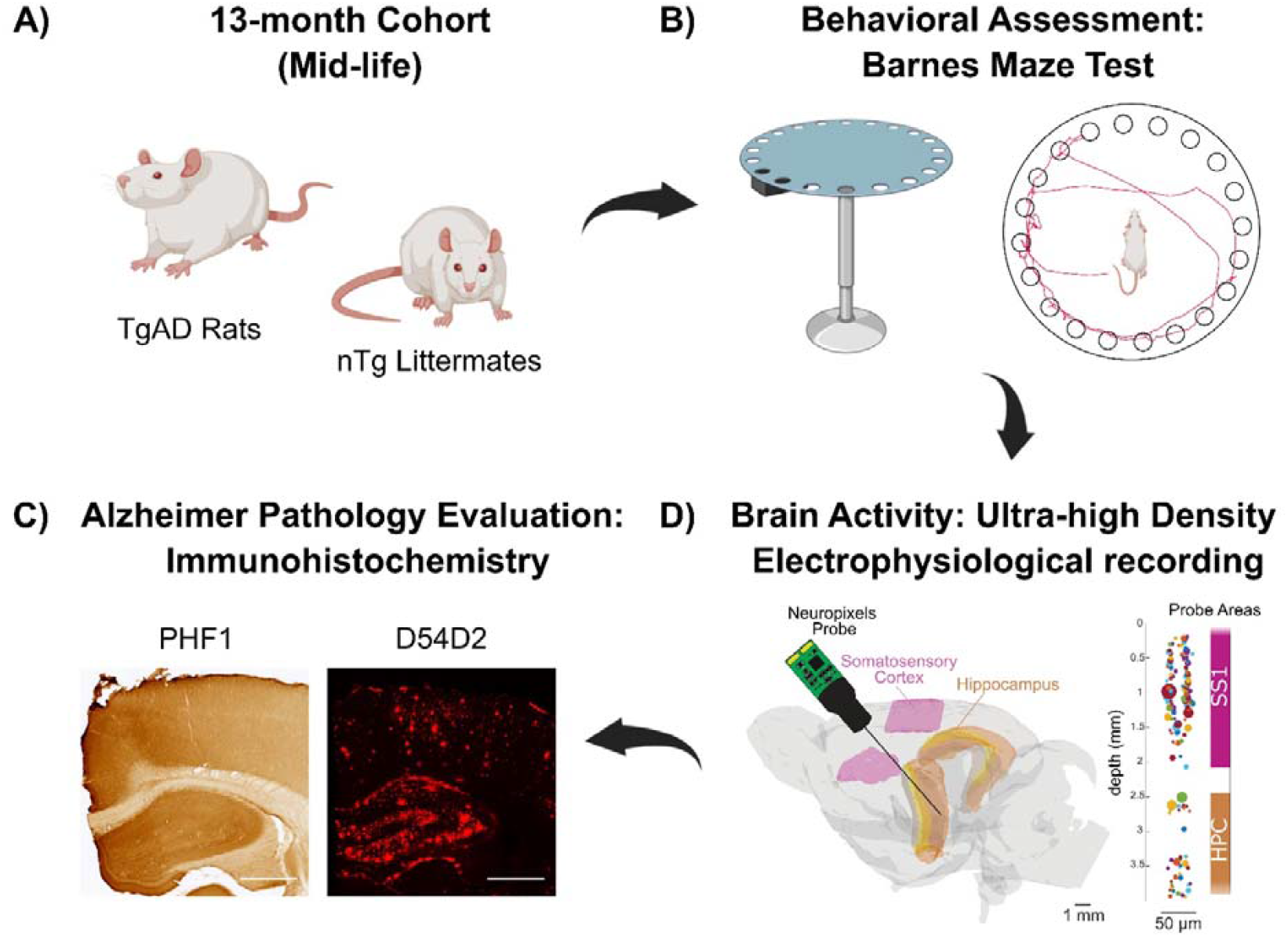
Experimental Design. A) Transgenic Fischer 344 rats overexpressing human APP with the Swedish mutation and PSEN1 mutation were used as a model of Alzheimer’s disease (AD). At 13 months of age, TgAD rats exhibit established amyloid and tau pathologies that reflect the early symptomatic phase of AD. Age-matched non-transgenic (nTg) littermates served as controls for normal aging. B) The rats’ spatial learning and memory were assessed using the Barnes maze test, where escape latency and search trajectories were recorded. C) Neuronal activity in the somatosensory cortex and hippocampus was recorded using high-density Neuropixels probes, allowing simultaneous detection of hundreds of single units. Forepaw stimulation at 3 Hz was applied to evoke brain responses. D). Immunohistochemistry for tau (PHF1) and Aβ (D54D2) was performed to assess pathological burden and test its association with cognitive resilience. White scale bar = 1mm.

### Identification of Cognitively Resilient TgAD Rats with established AD pathology

To assess cognitive outcomes, we conducted the Barnes Maze test in both TgAD and nTg rats at the symptomatic stage (13 months) and young adulthood (4 months), prior to amyloidogenesis (**Fig. 2A**). Age-related contrasts were apparent (**Fig. 2B**), especially with 13-month-old rats showing significantly slower reductions in escape latencies over reversal learning trials (p < 0.05). Moreover, we noticed that a subset of 13-month-old rats performed comparably to 4-month-old rats, and some aged rats showed high day-to-day variability which resembles the fluctuating cognitive performance often seen in early-stage dementia patients ^37,38^. Because individual differences are obscured by group-level averaging, we employed three area under the curve (AUC) metrics of escape latencies to better quantify the overall performances across individual rats’ learning and reversal learning trajectories: (1) AUC of escape latency across all learning trials (Learning trial 1-6), reflecting the rat’s ability to acquire and retain new spatial information; (2) AUC of latency during the first three days of reversal learning (reversal learning trial 1–6), assessing how rapidly the animal adapted to the new escape location; and (3) AUC of latency across the last two days of reversal learning (reversal learning trial 7–10), evaluating the consistency in cognitive flexibility and executive function (**Fig. 2C**). The variability of these 3 AUC parameters was elevated for 13-month-old TgAD and nTg rats, when compared to those of the young cohorts (Levene’s test; Learning t1–t6: p < 0.05; Reversal t1–t6: p < 0.05; Reversal t7–t10: p < 0.05).

**Figure 2:**
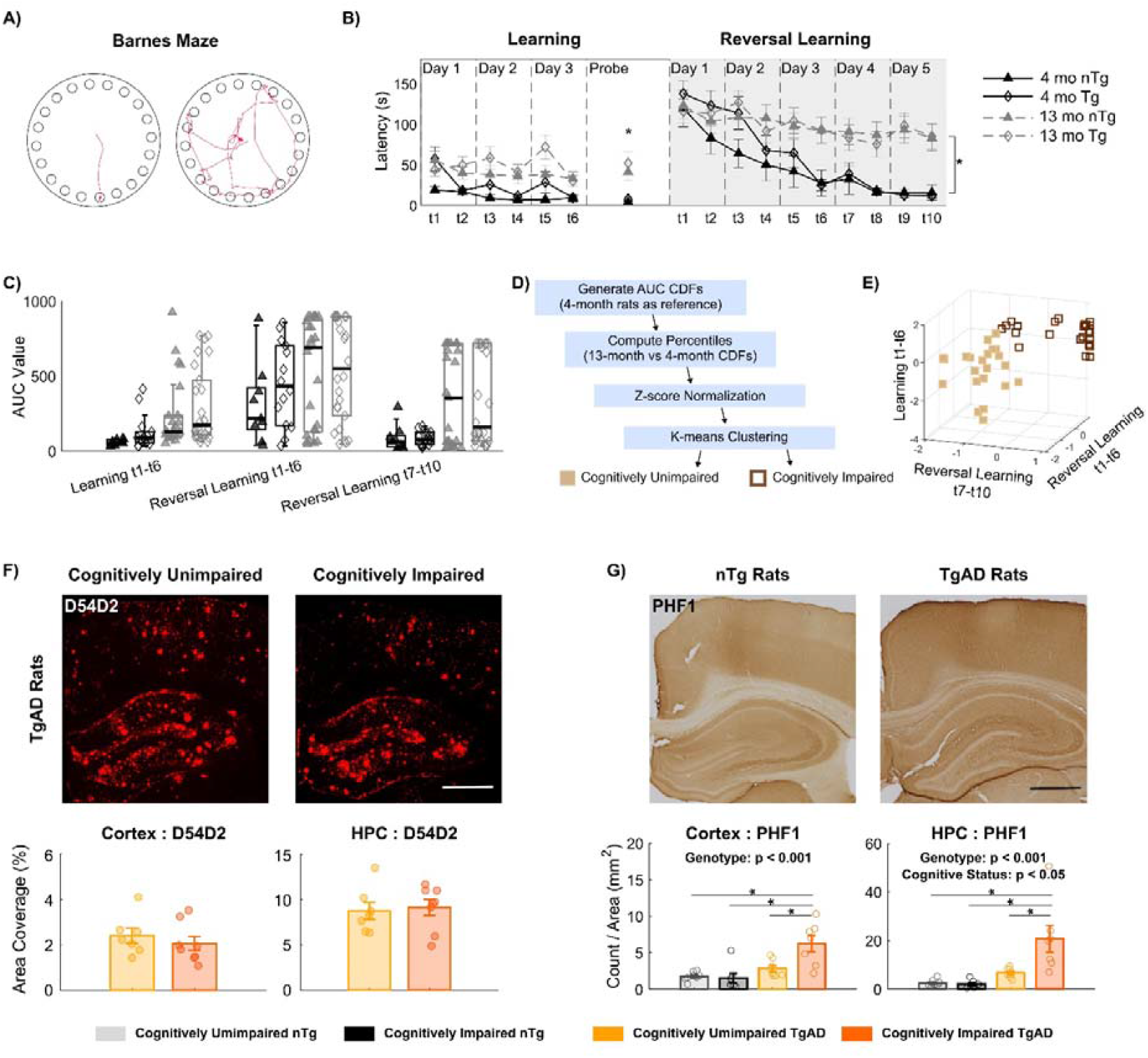
Maintained cognitive performance identifies resilient cohorts. **A)** The Barnes Maze test measures a rat’s ability to learn and remember the escape location in an aversive environment. Representative trajectories from trials with short and long escape latencies. **B)** This age-related decline in escape latencies was evident. 13-month-old rats showed significantly longer escape latency than did the 4-month-old rats in the spatial memory “probe” trial (p = 0.00289). While all rats showed a general reduction in escape latency over reversal trials, 13-month-old rats exhibited significantly less improvement than did the 4-month-old rats (p = 5.14e-04). **C)** High variability in cognitive performance among 13-month-old rats resulted in clear separation of AUCs in both TgAD and nTg cohorts. **D)** Schematic of the clustering pipeline used to determine cognitive outcomes based on the performance profiles of 4-month-old reference rats. **E)** K-means clustering of cognitive performance at 13 months identified 22 rats (10 nTg, 12 TgAD) as cognitively unimpaired and 31 rats (16 nTg, 15 TgAD) as cognitively impaired. **F)** β-amyloid burden quantified as the percent area coverage of D54D2 did not differ between cognitively unimpaired and impaired TgAD rats in the cortex (2.41 ± 0.86 vs. 2.07 ± 0.87) and HPC (8.75 ± 2.48 vs. 9.13 ± 2.46). **G)** PHF1 immunostaining revealed established tau pathology in 13-month-old TgAD rats versus nTg rats. Cognitively impaired TgAD rats (S1FL: 2.85 ± 1.19; hippocampus: 6.71 ± 2.09) had greater PHF1-positive inclusion density than did the cognitively unimpaired TgAD (S1FL: 6.26 ± 2.93; hippocampus: 20.74 ± 14.51). Scale bar = 1 mm. * denotes p < 0.05. Mean ± standard deviation.

To establish a reference for unimpaired cognition, we generated cumulative distribution functions (CDFs) for each AUC parameter using data from 4-month-old rats (n = 24, 12F, 12M). The performance of each 13-month-old rat was then compared against these 4-month-old CDFs to obtain percentile ranks. Finally, we applied k-means clustering to classify the aged rats into two groups (**Fig. 2D**). The cluster of rats whose performance was comparable to the young reference group was deemed as (**a**) “cognitively unimpaired” (nTg: 6M, 4F; Tg: 6M, 6F); whereas the other cluster, comprising rats with poorer performance across the AUC measures, was labeled as (**b**) “cognitively impaired” (nTg: 9M, 7F; Tg: 7M, 8F). To evaluate the AD pathology in these animals, we performed immunostaining for D54D2 (a marker for amyloid-β) ^39^ and PHF1 (a marker of pre-tangle tau) ^40^. The percent area covered by D54D2 was not significantly different between cognitively unimpaired and impaired TgAD rats in either the S1FL cortex or hippocampus (HPC) (**Fig. 2F**). Thus, we identified a cognitively resilient subset of 13-month-old TgAD rats that maintain cognitive performance despite their amyloid burden being indistinguishable from their impaired littermates.

Yet, the density of PHF1 inclusion for tau pathology demonstrated a genotype effect with TgAD rats exhibiting higher PHF1 burden than nTg rats in both S1FL and HPC (**Fig. 2F**; p < 0.05). Within TgAD rats, cognitively unimpaired rats showed a significantly lower PHF1 inclusion density compared to that of cognitively impaired rats (p < 0.05). Despite tau burden tracks cognitive status more closely than amyloid, it still explains only part of the variability in cognitive outcomes as people with similar tau loads can follow divergent cognitive trajectories ^41,42^. Our rat study supported this: D54D2 coverage and PHF1 density were not sufficient either separately or together to stratify cognitive status of 13-month-old TgAD rats (**Supplementary Fig. 1**). These findings underscore the need to integrate tau with additional markers to capture cognitive resilience in AD.

For brevity, we refer to cognitively unimpaired TgAD rats as ‘AD Resilient’; cognitively unimpaired nTg rats as ‘Healthy Aged’; cognitively impaired TgAD rats as ‘AD Impaired’; and cognitively impaired nTg rats as ‘Aging Impaired.’

### Populational Neural Activity: Preserved High-Frequency Cortical and Reduced Low-Frequency Hippocampal Oscillations Underlie AD Resilience

A Neuropixels probe 1.0 penetrated the primary forelimb somatosensory cortex (S1FL) and hippocampal subfields in the right hemisphere under light isofluorine. Forepaw stimulation was delivered using subdermal gold-disk electrodes (3 Hz, 5 mA, 333 µs pulse duration). Electrophysiological data were analyzed across spatial scales, beginning with population-level activity. We first quantified the power of local field potential (LFP) oscillations, which reflects population-level activity arising from synchronous transmembrane currents. We asked whether cognitively resilient animals exhibited differences in LFP power in S1FL and HPC (**Fig. 3A**). **Fig. 3B** shows time–frequency power spectra for S1FL and HPC, demonstrating stimulation-evoked modulation of spectral power.

**Figure 3:**
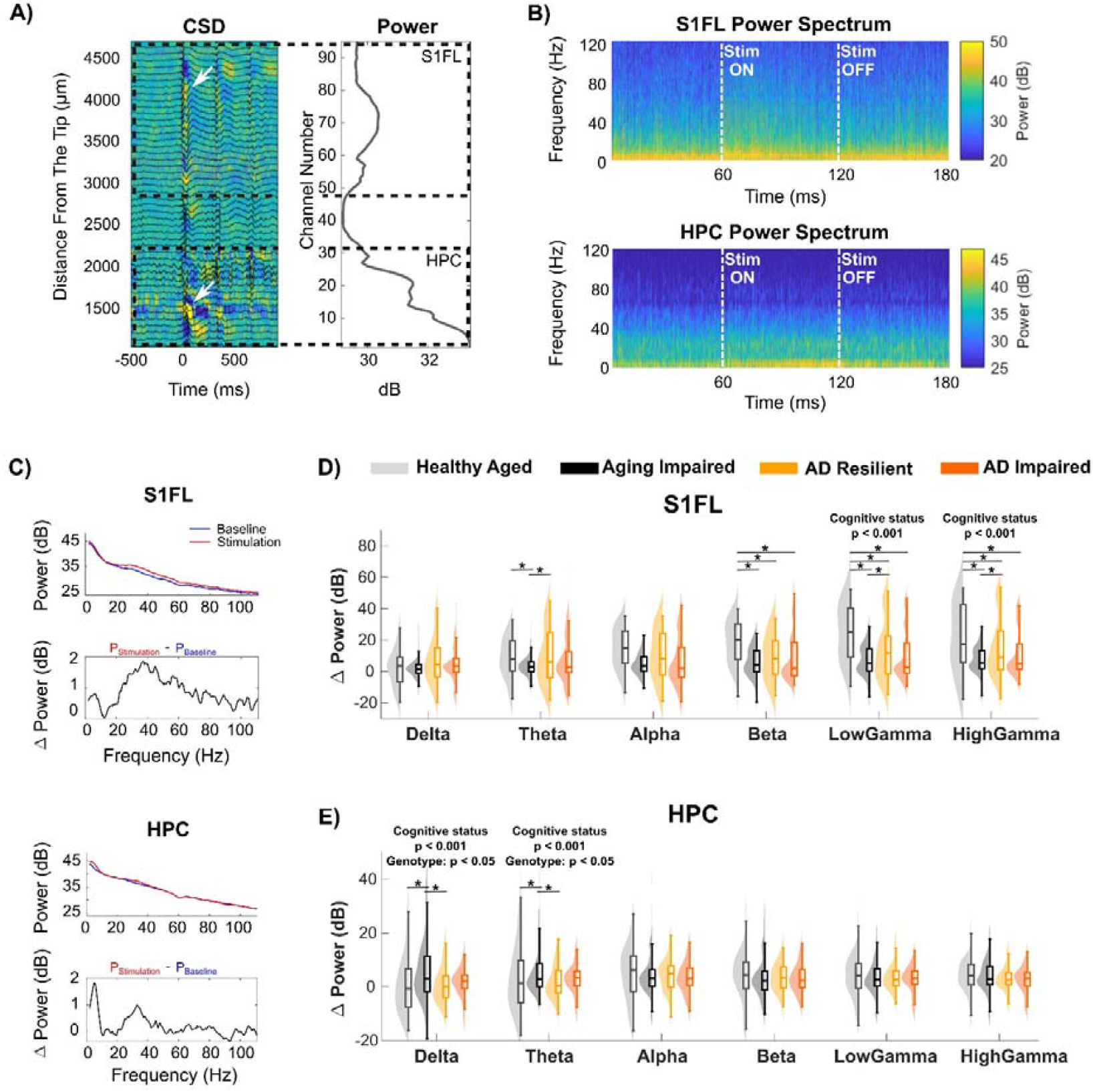
‘AD Resilient’ networks maintain evoked cortical gamma and hippocampal low-frequency oscillatory power. **A)** Current source density (CSD) map and 1–120 Hz local field potential (LFP) traces (left) and the corresponding spectral power distribution (right) along the inserted Neuropixels probe during 3 Hz forepaw stimulation. Strong sink–source couplings (indicated by white arrows) and power amplitude peaks were used to delineate the S1FL cortical and hippocampal regions. Yellow: sink; blue: source. **B)** Representative time–frequency power spectra for a 60-s forepaw stimulation trial averaged across channels in a representative cognitively unimpaired nTg Fischer 344 rat corresponding to cortical and hippocampal regions. **C)** Representative power spectra during baseline (blue) and forepaw stimulation (red) periods from single S1FL and HPC channels in an nTg rat depicted in (A). The normalized evoked power spectrum (*Δ Power* = *P*_*stimulation*_ - *P*_*baseline*_) revealed an elevated high-frequency oscillatory activity in S1FL and enhanced low-frequency oscillatory activity in the hippocampus. **D)** In the S1FL cortex, forepaw stimulation elicited significantly elevated low- and high-gamma power changes in ‘Healthy Aged’ and ‘AD Resilient’ rats when compared to those of cognitively impaired cohorts. Additionally, ‘Healthy Aged’ rats showed significantly higher evoked power compared to ‘AD Resilient’ rats (low gamma, p = 2.37e-06; high gamma, p = 0.0360). **F)** Evoked delta and theta power were significantly lower in ‘Healthy Aged’ and ‘AD Resilient’ rats relative to cognitively impaired groups, whereas TgAD rats exhibited higher delta/theta evoked power. * denotes p < 0.05.

The S1FL cortex exhibited enhanced evoked power predominantly in the gamma frequency range (30–80 Hz), whereas the HPC showed greater evoked power primarily within the low-frequency range (1–9 Hz) (**Fig. 3C**). In the S1FL, a significant main effect of cognitive status was observed in low-gamma and high-gamma frequency bands (**Fig. 3D**; p < 0.05). As gamma power reflects synchronized excitatory-inhibitory neural activity ^43–45^, ‘AD Resilient’ and ‘Healthy Aged’ rats showed significantly higher gamma evoked cortical power when compared to that of ‘AD Impaired’ and ‘Aging Impaired’ rats, suggesting preserved synchronized excitatory-inhibitory balance and network function that contribute to sensory perception.

In the HPC, a significant main effect of cognitive status was observed in the delta and theta bands (**Fig. 3E**). In contrast to the cortex, the HPC of ‘AD Resilient’ and ‘Healthy Aged’ rats exhibited significantly lower evoked power in these low-frequency bands (p < 0.05). Slow oscillations in delta and theta bands arise from excitatory drive that is paced by inhibitory networks ^46–48^, which are critical for spatial navigation and memory formation by the HPC^49,50^. The lower evoked theta and delta power observed in ‘AD Resilient’ and ‘Healthy Aged’ aged rats suggests well-balanced excitatory–inhibitory activity that supports task-relevant processing while avoiding excessive activation and the associated metabolic costs. Additionally, TgAD rats had significantly higher evoked power in the delta and theta bands compared to nTg rats^46–48^ (p < 0.05), suggesting that Alzheimer’s pathology disrupts normal oscillatory activities in the HPC. Together, these results indicate that “AD Resilient” rats, like “Healthy Aged” rats, have preserved cortical gamma activity and reduced low-frequency hippocampal activity.

### Fewer Activated Neurons but Elevated Evoked Firing Rate Underlie AD Resilience

Having observed the frequency-specific, region-dependent differences in population-level LFP power, we next analyzed single-unit recordings to determine how the activity of individual neurons differs between resilient and non-resilient rats. The use of Neuropixels probes allowed detection of hundreds of single units per animal. Single units with somatic waveforms, most presumptively located within a 50 µm radius of the probe, were manually curated using Phy (**Fig. 4A**). These somatic single units were further classified as 5529 putative excitatory single units (hereafter, ‘excitatory neurons’) and 2815 putative inhibitory single units (hereafter, ‘inhibitory neurons’) based on their trough-to-peak latency (**Fig. 4B**). The signal quality of recorded single units including signal-to-noise ratio, peak-to-peak amplitude, and noise floor was characterized in **Supplementary Fig. 2**.

**Figure 4.**
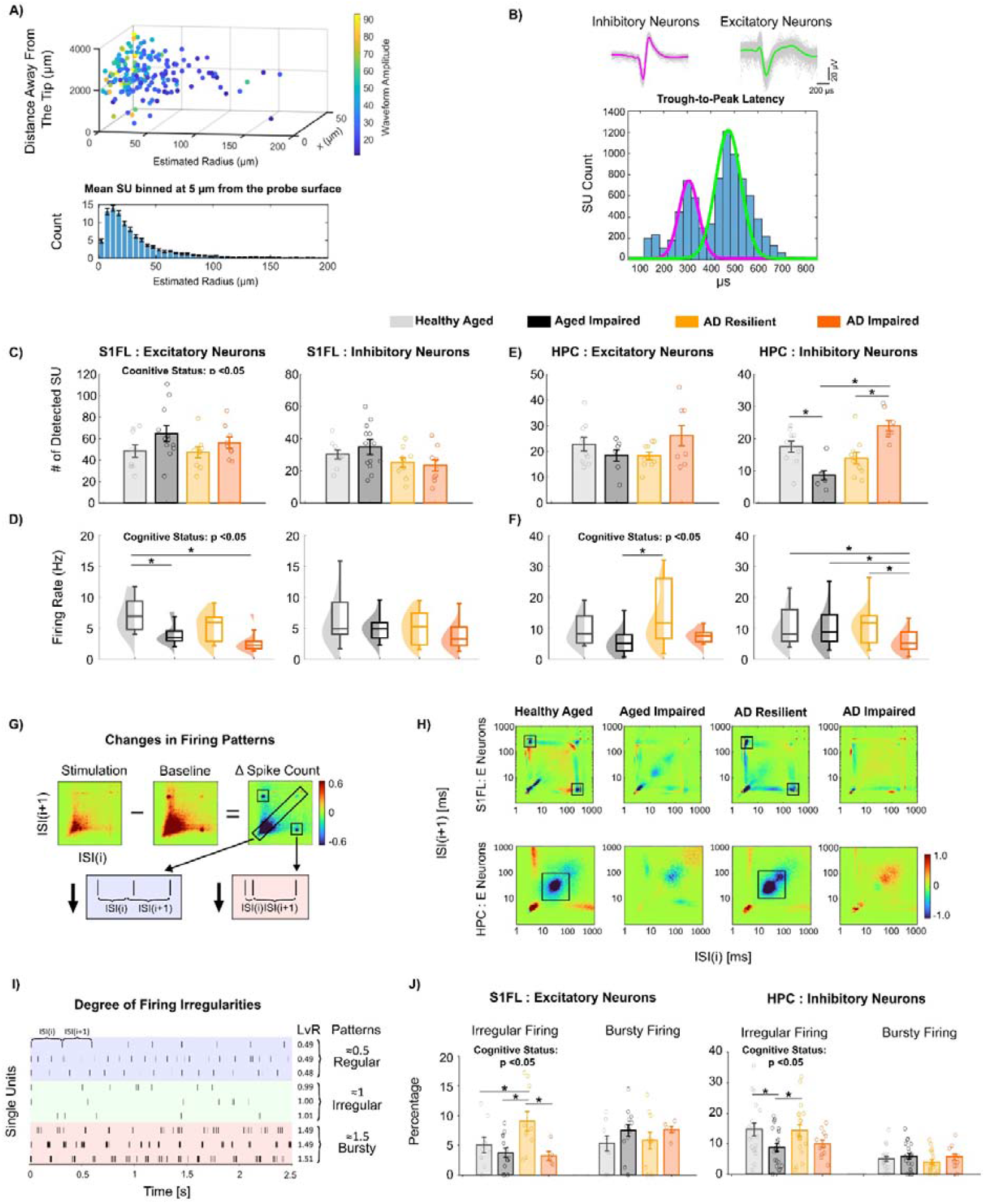
‘AD Resilient’ networks have fewer activated neurons with robust evoked firing and show reduced excitatory spike burstiness with increased irregular firing during forepaw stimulation. **A)** Estimated localization of sorted single units (SU) in 3D space, color-coded by their peak-to-peak amplitude. The x-axis represents the electrode position, y indicates the estimated radial distance of the SU from the electrode, and z denotes the distance from the probe tip. The projection onto the y-axis revealed that approximately 83.77% SUs were located within ~50 µm of the electrode surface. **B)**. Top: Example of a representative narrow waveform classified as a putative inhibitory neuron (left) and representative wide waveform sorted as a putative excitatory neuron (right). Bottom: Single-units (SUs) were classified based on trough-to-peak (TP) latency, which followed a bimodal distribution. SUs with TP latencies less than 0.38 ms were classified as putative inhibitory neurons (magenta), while SUs with TP latencies greater than 0.38 ms were classified as putative excitatory neurons (green). **C)** In S1FL, ‘Healthy Aged and ‘AD Resilient’ rats showed fewer activated excitatory neurons relative to ‘Aging Impaired’ and ‘AD Impaired’ groups. The number and evoked firing rates of activated S1FL inhibitory neurons remained comparable. **D)** The evoked firing rates of S1FL excitatory neurons during forepaw stimulation were significantly higher in ‘Healthy Aged’ and ‘AD Resilient’ rats compared to ‘Aging Impaired’ and ‘AD Impaired’ groups. **E)** The number of activated HPC inhibitory neurons was similar between ‘Healthy Aged’ (17.50±5.44) and ‘AD Resilient’ (14.56±5.73) rats, whereas ‘Aging Impaired’ (9.28±4.42) rats showed a significantly lower count, and ‘AD Impaired’ (24.00±4.90) rats exhibited a significantly higher count. **F)** HPC excitatory neurons in ‘AD Resilient’ and ‘Healthy Aged’ rats had significantly higher evoked firing rates than those in ‘Aging Impaired’ and ‘AD Impaired’ rats. Additionally, ‘AD Impaired’ (5.92±4.23) rats exhibited a significantly lower firing rate of putative inhibitory neurons relative to other groups (Healthy Aged: 11.36±7.27, Aging Impaired: 10.40±7.07; AD Resilient: 12.41±7.75). **G)** Stimulation-induced ISI changes were computed as (sensory stimulation − baseline). Patterns in ΔISI histograms can provide insight into the temporal spiking patterns of neurons: clusters in the short–long (*ISI*_*i*_ << *ISI*_*i* +1_) or long–short (*ISI*_*i*_ >> *ISI*_*i*+1_) corners suggest burstiness, whereas clusters along the main diagonal (*ISI*_*i* ≈_*ISI*_*i*+1_) indicate periodic spiking. Positive ΔISI bins indicate ISI ranges that occur more often during stimulation than baseline, and negative bins indicate reductions. **H)** ‘AD Resilient’ and ‘Healthy Aged’ rats exhibited reduced excitatory spike burstiness with distinct region-specific patterns in S1FL and HPC, patterns that were not observed in ‘AD Impaired’ or ‘Aging Impaired’ rats. **I)** Representative spike sequence patterns illustrating three firing regimes derived from Local Variation with Refractoriness correction (LvR) values centered around 0.5, 1.0, and 1.5, corresponding respectively to regular, irregular, and bursty dynamics. **J**) ‘AD Resilient’ and ‘Healthy Aged’ rats had significantly higher percentages of S1FL excitatory neurons and HPC inhibitory neurons with irregular firing patterns compared to ‘AD Impaired’ or ‘Aging Impaired’ rats.

We next asked how neurons responded during forepaw sensory stimulation by quantifying the number of activated neurons and their firing rates. During transcutaneous forepaw stimulation, cognitive status had a significant main effect on S1FL evoked firing rates of *excitatory* neurons: ‘AD Resilient’ and ‘Healthy Aged’ rats had significantly *fewer* detected excitatory neurons (**Fig. 4C**; p < 0.05) but significantly *higher* evoked firing rates than ‘AD Impaired’ and ‘Aging Impaired’ rats (**Fig. 4D**; p < 0.05). This observation suggests that in ‘AD Resilient’ and ‘Healthy Aged’ rats a smaller number of excitatory neurons is preferentially and more strongly involved during S1FL network activation. In the HPC, ‘AD Resilient’ and ‘Healthy Aged’ rats exhibited significantly *higher* evoked excitatory firing rates than ‘AD Impaired’ and ‘Aging Impaired’ rats (**Fig. 4F**; p < 0.05). In HPC inhibitory neurons, the effects of cognitive status diverged between aged nTg vs TgAD cohorts (**Fig. 4E-F**), ‘Healthy Aged’ rats exhibited significantly more activated inhibitory neurons than ‘Aging impaired rats’ (p < 0.05). However, the ‘AD Resilient’ showed significantly fewer activated inhibitory neurons (p < 0.05), but significantly higher evoked inhibitory firing rates (p < 0.05) compared to ‘AD impaired’ rats. The immunostaining revealed no significant group difference in the cell densities of either GAD67+NeuN+ inhibitory neurons or GAD67-NeuN+ excitatory neurons in S1FL cortex, hippocampal CA1, and CA3 regions, indicating that the observed differences in electrophysiological SU profiles reflect functional changes rather than variations in neuronal population size (**Supplementary Fig. 3B**). Together, these results suggest that cognitive resilience in AD pathology manifests as fewer activating neurons coupled with elevated evoked firing rates, particularly in cortical excitatory and HPC inhibitory populations.

### Reduced Excitatory Spike Burstiness and Increased Irregular Firing Neurons Underlies AD Resilience

Given the effect of cognitive status on excitatory firing rates in both S1FL and HPC, we next examined whether spike temporal dynamics distinguish cognitive resilience from impaired phenotypes. To determine how forepaw-evoked network activation alters spike-timing dynamics, we quantified the changes in inter-spike interval (ISI) between forepaw stimulation and baseline periods (ΔISI; **Fig. 4G**). In S1FL, excitatory neurons in “AD Resilient” and “Healthy Aged” rats showed reduced burstiness in the short–long ISI ROI (~3-ms followed by ~200-ms ISI pairs; **Fig. 4H**, boxed ROI). “Healthy Aged” rats exhibited nearly a 1.8-fold greater reduction in spike-count than that of “Aging Impaired” rats; similarly, “AD Resilient” rats showed approximately a 5-fold greater reduction than did “AD Impaired” rats. In the HPC, excitatory neurons in the cognitively unimpaired groups showed reduced periodic spiking at ~50 ms (**Fig. 4H**). “Healthy Aged” rats exhibited an approximately 7-fold greater reduction than “Aging Impaired” rats, and “AD Resilient” rats showed a nearly 5.6-fold greater reduction than “AD Impaired” rats. In contrast to excitatory neurons, inhibitory cells showed no differences in their ΔISI histograms in either S1FL or HPC (**Supplementary Fig. 4A**). Together, these results suggest that cognitive resilience is associated with region-specific reductions of excitatory burstiness during network activation.

To examine whether spike-timing patterns differ by genotype and cognitive status, we quantified ISI variability using the Local Variation with Refractoriness correction (LvR; **Supplementary Fig. 4B**). LvR values (**Fig. 4I**) characterize spike train dynamics as regular (~0.5), irregular(~1.0), or bursty (~1.5), independent of firing rate and spike counts ^51^. “AD Resilient” and “Healthy Aged” rats had a significantly higher proportion of S1FL excitatory neurons (p < 0.05) and HPC inhibitory neurons (p < 0.05) with irregular firing compared with ‘AD Impaired’ and ‘Aging Impaired’ rats (**Fig. 4J**). Furthermore, we observed a significant genotype effect for S1FL inhibitory neurons, with TgAD rats exhibiting a greater proportion of irregular-firing S1FL inhibitory neurons compared to nTg impaired rats (**Supplementary Fig. 4C**; p < 0.05). Collectively, these observations indicate that in cognitively resilient rats, cortical excitatory and HPC inhibitory neurons fire more irregularly than they do in cognitively impaired rats

### Region- and Subtype-Specific Stability of Neuronal Representations at Population-Level Supports AD Resilience

Although distinct patterns in neuronal activity were identified across cognitive status and genotype, measures such as firing rate are averaged across trials and obscure variability in neuronal responses to repeated stimulation. Despite documented variability in individual neuronal responses, the overlapping neuron population supports efficient information processing and adaptive plasticity in normal brain function ^30^. Therefore, we next asked whether reliable neuronal representations during repeated stimulation were associated with cognitive resilience. To quantify this, we defined the neuronal engagement index (NEI) as the proportion of the trial’s stimuli that evoked a significant change in the neuron’s firing rate relative to that at baseline (permutation test, p < 0.05). Meanwhile, this NEI index mathematically represents the proportion of neurons activated by a given stimulus relative to the total number of neurons that could be activated within a trial (see Methods). A higher NEI value indicates more consistent recruitment of the same neuronal subset across stimulus pulses, reflecting more stable neuronal representations (**Fig. 5A**). A significant main effect of cognitive status was observed in the S1FL excitatory neurons (**Fig. 5B**; p < 0.05) and hippocampal inhibitory neurons (p < 0.05). In both neuronal populations, ‘AD Resilient’ and ‘Healthy Aged’ rats exhibited significantly higher NEI values compared to ‘AD Impaired’ and ‘Aging Impaired’ rats, suggesting a greater stability of neuronal representations. No significant differences were observed in the average number of neuronal subsets activated per stimulation pulse. However, there was a trend toward group differences in the total number of activated neurons across all stimulation pulses within a single trial for both excitatory and inhibitory neurons in S1FL and HPC (**Supplementary Fig. 5**), consistent with the significant cohort differences in the total number of activated neurons across all trials of each animal shown in **Fig. 4A, 4C**. Moreover, we observed a significant inverse relationship between cognitive performance and NEI in S1FL excitatory neurons and hippocampal inhibitory neurons, indicating higher NEI values were associated with lower latency AUCs particularly in the reversal learning phase (**Fig. 5C; Supplementary Fig. 6**). In summary, these results indicate that stable population-level representations of cortical excitatory and hippocampal inhibitory neurons during repeated stimulation underlie resilience to cognitive decline associated with Alzheimer’s pathology and aging.

**Figure 5:**
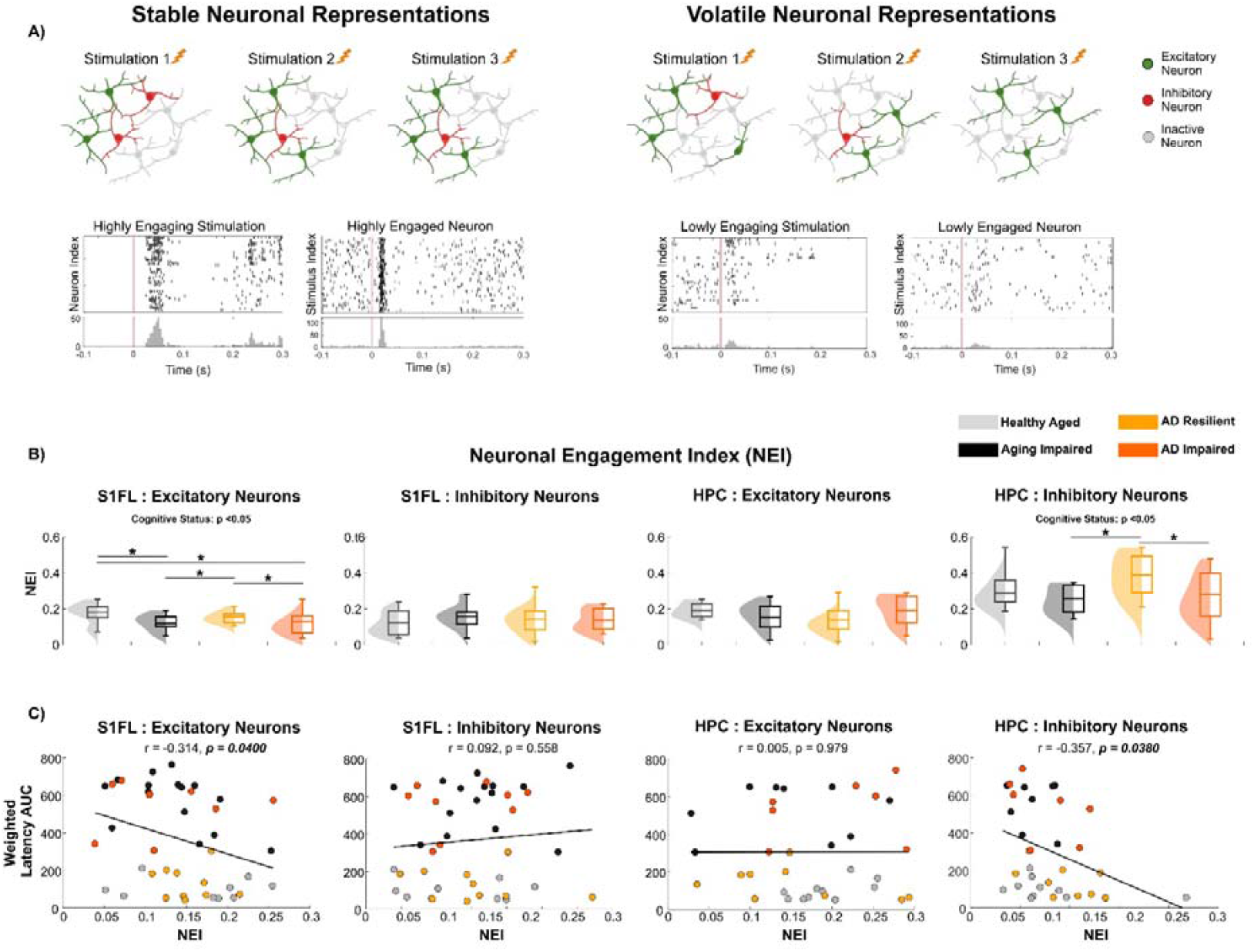
‘AD Resilient’ rats exhibit stable population-level representations. **A)**. Schematic illustration of the concept of neuronal representations during repeated forepaw stimulation. In networks with stable population-level representations, a consistent subset of neurons is repeatedly activated across trials, indicating reliable and overlapping neuronal engagement. In contrast, volatile representations involve a lower overlap of activated neurons, where different neuronal subsets respond across trials, reflecting less consistent activation patterns and reduced response reliability. **B)** ‘Healthy Aged’ and ‘AD Resilient’ rats showed elevated Neuronal Engagement Index (NEI) compared to ‘Aging Impaired’ and ‘AD Impaired’ rats for S1FL excitatory neurons and HPC inhibitory neurons. **C)** Pearson’s correlation coefficients between the weighted composite escape-latency AUC score (0.2·Learning t1–t6 AUC + 0.3·Reversal Learning t1–t6 AUC + 0.5·Reversal Learning t7–t10 AUC) and NEI were significant for S1FL excitatory neurons and hippocampal inhibitory neurons.

### Preserved Inhibitory Synaptic Connections Underlie AD Resilience

As the pattern of synaptic connectivity shapes how often a neuron is recruited during network activation, we next investigated whether patterns in synaptic connections could explain the reliable neuronal representation associated with preserved cognition. Cross-correlograms (CCG) of simultaneously recorded single units were used to investigate monosynaptic connectivity patterns.

In the S1FL, the number of CCG-identified monosynaptic connections were comparable across groups (**Fig. 6C**). Yet, ‘AD Resilient’ and ‘Healthy Aged’ rats had a significantly lower percentage of CCG-based excitatory connections and a correspondingly higher percentage of inhibitory connections relative to ‘AD Impaired’ and ‘Aging Impaired’ rats (**Fig. 6D**; p < 0.05). For CCG-based excitatory connections, neither connection strength (**Supplementary Fig. 7A**) nor the proportion of E→E versus E→I connections differed significantly across groups (**Fig. 6E**). In contrast, ‘AD Resilient’ and ‘Healthy Aged’ rats exhibited a significantly higher CCG-based inhibitory connection strength (**Fig. 6F**; p < 0.05) as well as a significantly higher proportion of I→E and a lower proportion of I→I connections (**Fig. 6G**; p < 0.05) than ‘AD Impaired’ and ‘Aging Impaired’ rats. These findings suggest that cognitive resilience in AD pathology in S1FL manifests as fewer excitatory synaptic connections and more stronger inhibitory synaptic inputs targeting excitatory neurons, supporting the maintenance of gamma-band LFP in S1FL (**Fig. 3**).

**Figure 6:**
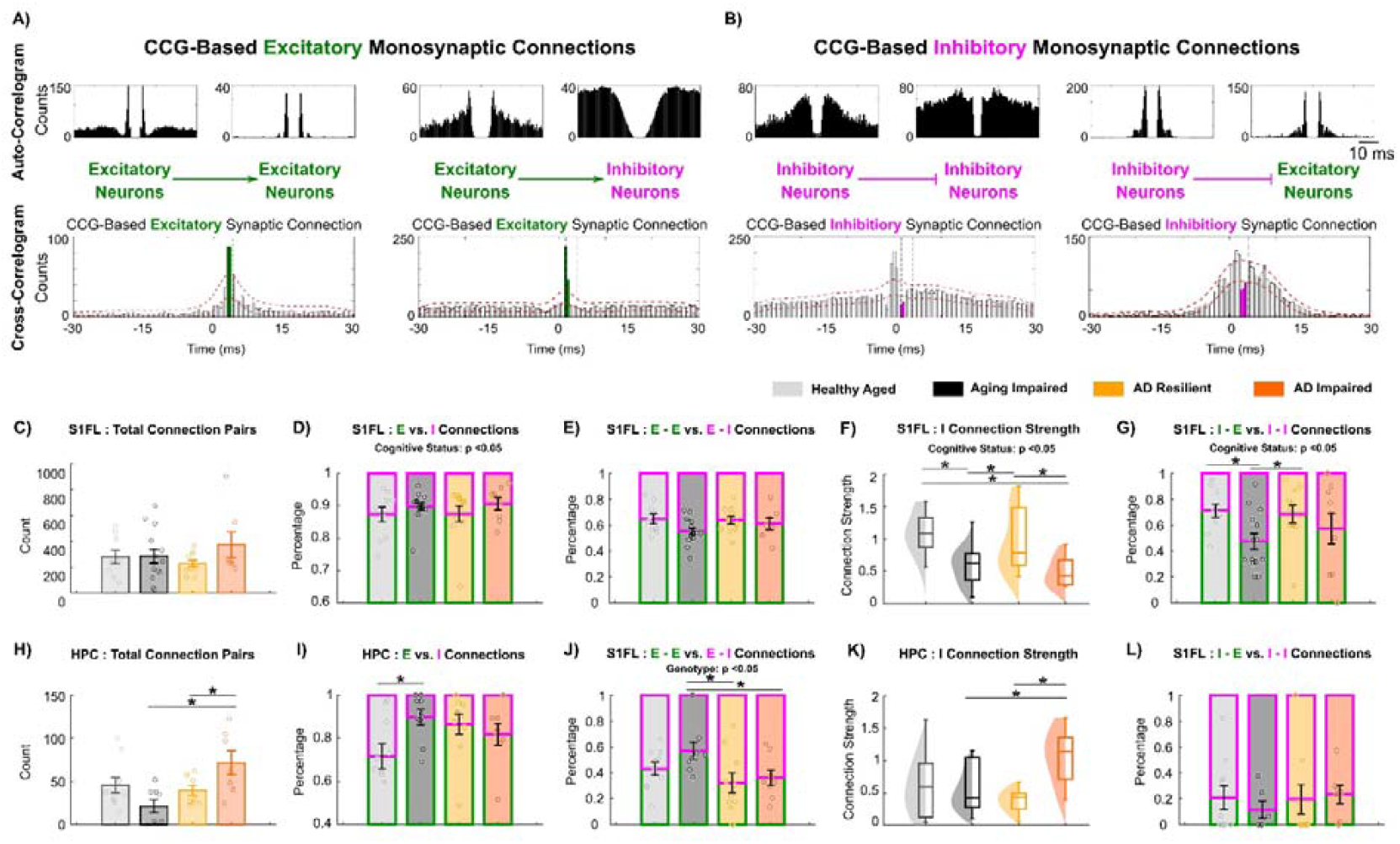
AD Resilient networks maintain putative excitatory and inhibitory synaptic connections. **A)** Representative cross-correlogram (CCG) illustrating a monosynaptic excitatory-to-excitatory (E-E) and a monosynaptic excitatory-to-inhibitory (E-I) connection. Autocorrelograms of each neuron are shown above. Excitatory synaptic connection was identified by a short-latency (<4 ms) peak in postsynaptic spiking that exceeded the red dotted significance threshold. Waveform-based classification was further used to validate the CCG-identified presynaptic units. **B)** Representative CCG showing a monosynaptic inhibitory-to-inhibitory (I-I) and a monosynaptic inhibitory-to-excitatory (I-E) connection. Inhibitory synaptic transmission was inferred from a short-latency (<4 ms) suppression of postsynaptic firing below the red dotted significance threshold. Neuron subtypes were classified by trough-to-peak (TP) latency as described in Figure 4. **C)** The total number of CCG-identified monosynaptic connections in S1FL was comparable across genotypes and cognitive groups in the S1FL cortex (Genotype: p = 0.486; Cognitive Status: p = 0.891). **D)** In S1FL, the percentages of excitatory (E) and inhibitory (I) connections onto postsynaptic partners revealed a significant effect of cognitive status (p = 0.0419), with ‘AD Resilient’ (12.448.16%) and ‘Healthy Aged’ (13.546.82%) rats exhibiting a higher relative proportion of inhibitory compared to ‘Aging Impaired’ (10.184.51%) and ‘AD Impaired’ (7.714.97%) rats. **E)** The percentage of E-E (and corresponding E-I) connections in S1FL were statistically comparable across groups (Genotype: p = 0.467; Cognitive Status: p = 0.0665). **F)** In the S1FL cortex, ‘AD Resilient’ (0.980.50) and ‘Healthy Aged’ (1.100.33) rats exhibited significantly stronger inhibitory synaptic connection strength (p = 9.53e-05) than did the ‘Aging Impaired’ (0.590.31) and ‘AD Impaired’ (0.500.25) rats. **G)** In the S1FL cortex, the percentages of I-E vs. I-I connections relative to the total number of CCG-identified inhibitory transmissions revealed that ‘AD Resilient’ (68.48 ± 22.40%) and ‘Healthy Aged’ (71.21 ± 16.37%) rats had a significantly lower proportion of I→E connections (p = 0.00907) compared to’Aging Impaired’ (47.45 ± 22.33%) and ‘AD Impaired’ (58.12 ± 37.84%) rats. **H)** In HPC, the total number of CCG-based synaptic connections was significantly lower in ‘AD Resilient’ rats than those in ‘AD Impaired’ rats (p = 0.0130). **I)** ‘Healthy Aged’ rats exhibited a significantly higher percentage of CCG-based inhibitory (28.2118.14%) connections in HPC compared to the other three groups. **J)** In HPC, a significant genotype effect was observed with nTg rats (Healthy Aged: 43.20 ± 15.54%, Aging Impaired: 53.79 ± 29.52%) exhibiting a significantly higher E-E to E-I ratio than nTg rats (AD Resilient: 35.91 ± 22.24%, AD Impaired: 36.64 ± 18.01%). **K)** In the hippocampus, only ‘AD Impaired’ rats (1.060.44) exhibited significantly elevated CCG-based inhibitory synaptic connection strength (vs. Healthy Aged: 0.670.53, Aging Impaired: 0.610.45, AD Resilient: 0.380.19). **L)** In the HPC, I→I connections constitute the majority of inhibitory synapses across groups. * denotes p < 0.05. Mean ± standard deviation.

In the HPC, the effects of cognitive status diverged between aging and AD cohorts. ‘AD Resilient’ rats had significantly fewer CCG-based monosynaptic connections and a trend of a lower percentage of inhibitory synaptic connections than did ‘AD impaired’ rats (**Fig. 6H**; p < 0.05); instead, ‘Healthy Aged’ rats showed a significantly *higher* percentage of inhibitory connections (**Fig. 6I**; p < 0.05) than ‘Aging Impaired’ rats. These observations suggested that resilience to aging-dependent cognitive decline is associated with greater hippocampal inhibitory synaptic connections. TgAD rats displayed a significantly lower percentage of E→E connections and a correspondingly higher percentage of E→I connections than did nTg rats (**Fig. 6J**; p < 0.05), indicating AD pathologies shift HPC excitatory synaptic inputs toward inhibitory neurons. “AD Impaired” rats also showed significantly higher HPC inhibitory synaptic strength than did ‘AD Resilient’ rats (**Fig. 6K**; p < 0.05), and HPC I→I connections predominated across all four groups (**Fig. 6L**).

Collectively, cognitive resilience in AD pathologies in HPC manifests as the prevention of excessively strong I→I synaptic connections in the context of an excitatory drive biased toward inhibitory neurons, likely constraining disinhibition and, in turn, suppressing network hyperexcitation, consistent with preserved low-frequency LFP power shown in **Fig. 3**.

We also found that the intersomatic distance in CCG-identified pairs was comparable across groups (**Supplementary Fig. 7B**), suggesting that the observed shifts in the proportion of CCG-based excitatory vs. inhibitory synaptic connections and in inhibitory connection strength are likely driven by functional reorganization rather than morphological changes.

### Constrained Excitatory Synaptic Convergence and Inhibitory Synaptic Divergence onto Inhibitory Neurons Underlies AD Resilience

As a neuron can receive inputs from many presynaptic partners, we quantified excitatory convergence and inhibitory divergence to determine whether microcircuit organisation accounts for cognitive resilience (**Fig. 7A, 7D**). In S1FL, ‘AD Resilient’ and ‘Healthy Aged’ showed significantly *fewer* excitatory neurons receiving >5 convergent excitatory inputs than ‘AD Impaired’ and ‘Aging Impaired’ rats (**Fig. 7B**; p < 0.05). ‘Healthy Aged’ rats showed significantly *more* inhibitory neurons diverging to 2–5 excitatory targets than did ‘Aging Impaired’ rats (**Fig. 7E**; p < 0.05). Yet, ‘AD Resilient’ and ‘Healthy Aged’ had significantly fewer inhibitory neurons diverging to 2-5 inhibitory neurons than ‘AD Impaired’ and ‘Aging Impaired’ rats (**Fig. 7F**; p < 0.05). Thus, cognitive resilience in S1FL manifests as reduced excitatory convergence onto excitatory cells, greater inhibitory output to excitatory neurons, and restricted inhibitory– to–inhibitory divergence.

**Figure 7:**
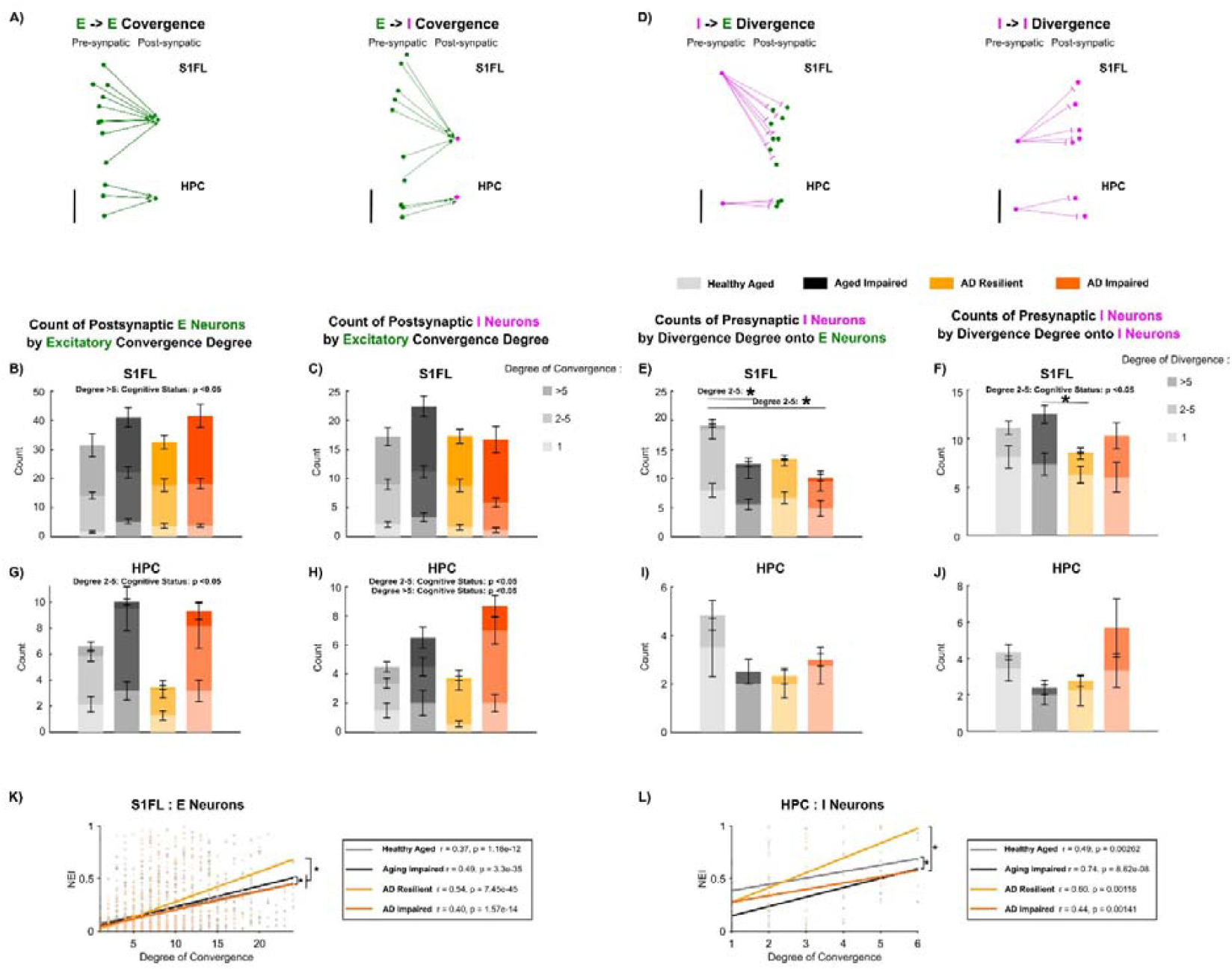
‘AD Resilient’ networks exhibit distinct synaptic connectivity patterns that correlate to stable neuronal representations. **A)** Representative examples of excitatory-to-excitatory (E-E) (left) and excitatory-to-inhibitory convergence (right) in the S1FL and hippocampus. The degree of convergence measures the number of presynaptic excitatory neurons forming monosynaptic inputs onto a single postsynaptic excitatory or inhibitory neuron. In S1FL, the ‘Healthy Aged’ rats tended to have fewer excitatory neurons **B)** and inhibitory neurons **C)** that received >1 excitatory inputs. **D)** Representative examples of inhibitory-to-excitatory (I-E) and inhibitory-to-inhibitory (I-I) divergence in the S1FL cortex and hippocampus (left). The degree of divergence quantifies the number of postsynaptic excitatory neurons connected with a single presynaptic inhibitory neuron. **E)** In S1FL, ‘Healthy Aged’ rats had significantly more inhibitory neurons that diverged onto 2-5 excitatory neurons than other groups **F)** Analysis of S1FL inhibitory-to-inhibitory (I→I) divergence revealed that ‘AD Resilient’ and ‘Healthy Aged’ rats had significantly fewer presynaptic inhibitory neurons forming connections with 2–5 postsynaptic inhibitory partners compared to other groups. In HPC, ‘AD Resilient’ and ‘Healthy Aged’ rats showed significantly fewer postsynaptic excitatory **G)** and inhibitory **H)** neurons receiving 2–5 or >5 convergent excitatory inputs relative to cognitively impaired rats. I) ‘Healthy Aged’ tended to have more HPC inhibitory neurons that diverged onto 2-5 postsynaptic excitatory neurons than other three groups. J) ‘AD resilient’ rats tended to have fewer HPC inhibitory neurons that diverged onto 2-5 inhibitory neurons than ‘AD Impaired’ rats. **K)** Pearson’s correlation coefficient between the degree of convergence of postsynaptic putative excitatory neurons in the S1FL cortex and their Neuronal Engagement Index (NEI) was significantly higher in the ‘AD Resilient’ rats than in the other 3 groups. **L)** Pearson’s correlation coefficient between the degree of convergence of postsynaptic putative inhibitory neurons in the hippocampus and their Neuronal Engagement Index (NEI) was significantly higher in ‘AD Resilient’ rats than in the other groups. Scale bar = 500*µ*m. * denotes p < 0.05. Mean ± standard deviation.

In the HPC, ‘AD Resilient’ and ‘Healthy Aged’ rats had significantly *fewer* excitatory neurons receiving 2–5 excitatory inputs (**Fig. 7G**; p < 0.05) and significantly *fewer* inhibitory neurons receiving 2-5 excitatory inputs (**Fig. 7H**; p < 0.05) compared to ‘Aging Impaired’ and ‘AD Impaired’ rats. ‘AD Resilient’ rats also showed a trend toward fewer inhibitory neurons diverging to 2–5 inhibitory targets compared with ‘AD Impaired’ rats (p = 0.142; **Fig. 7J**). Together, these data indicate that AD resilience in HPC is associated with reduced excitatory convergence onto both excitatory and inhibitory neurons and fewer inhibitory neurons targeting multiple inhibitory neurons, complementing the constrained I→I coupling in Fig. 6.

We next linked excitatory convergence to the reliability of neuronal representations (NEI). ‘Healthy Aged’ rats had significantly lower correlation between the degree of convergent excitatory input and NEI in S1FL excitatory neurons (**Fig, 7K**; p < 0.05) and inhibitory neurons (**Supplementary Fig. 8A**; p < 0.05), relative to ‘Aging Impaired’ rats. In contrast, with AD pathology, ‘AD Resilient’ rats showed greater excitatory convergence onto S1FL excitatory neurons (**Fig, 7K**; p < 0.05) and HPC inhibitory neurons (**Fig, 7L**; p < 0.05) accompanied by significantly higher NEI than that in ‘AD Impaired’ rats. These data suggest that reliance of neuronal representation stability on excitatory convergence is context dependent: weaker in cognitive resilience in non-diseased aging, but stronger in cognitive resilience in AD pathologies.

## Discussion

Amyloid and tau proteinopathies are thought to be drivers of cognitive impairment in AD ^52–54^, yet a plethora of pharmacological agents targeting these proteins have had modest impact on cognitive decline ^55–57^. Research interest has therefore shifted toward cognitive resilience, as clarifying its mechanisms may offer a novel therapeutic opportunity to prevent or delay dementia in susceptible populations. Here, we identified stable population-level representations of cortical excitatory and hippocampal inhibitory neurons as a hallmark of cognitive resilience in the presence of AD pathology. We further demonstrated that cognitive resilience is associated with region-specific distinctions in functional synaptic connectivity that may help re-establish excitation–inhibition balance despite accumulating amyloid and tau pathologies. Our findings highlight a potential biomarker to guide neuromodulation strategies for preserving brain function in aging and neurodegeneration.

### Excitatory circuits – S1FL

In both AD resilience and healthy aging, S1FL activation during repeated sensory stimulation is supported by a smaller pool of excitatory neurons and fewer excitatory synaptic connections than in Aging-Impaired and AD-Impaired groups (**Figs. 4C, 6D, 7B**). This selective recruitment may reduce the metabolic cost of spiking under age-related constraints in mitochondrial and vascular functions ^30,58–60^. Encoding stimuli with fewer, consistently engaged neurons lowers energy expenditure while preserving accuracy. Selective and stable population coding may be key for sustaining cognitive function during aging and in AD. By contrast, in aging impairment and AD dementia, a larger population of cortical excitatory neurons is recruited, yet these neurons fire at lower rates and form unstable population-level representations during repeated stimulation (**Figs. 4A,B, 5**). Excitatory drive is upregulated, with increased excitatory-to-excitatory connectivity and greater convergence of excitatory inputs onto excitatory neurons (**Figs. 6D, 7B**), consistent with neuronal hyperexcitability reported in AD dementia ^61^. This combination of broadened, inefficient recruitment and strengthened recurrent excitation likely contributes to breakdown of cortical E/I balance in AD dementia.

### Excitatory circuits – HPC

Excitatory circuits in HPC also distinguish resilience, AD dementia, and aging impairment. In AD resilience, excitatory drive is redistributed away from excitatory neurons toward inhibitory circuitry relative to healthy aging. The distribution of synaptic inputs shifts toward greater overall excitatory drive (**Fig. 6I**), but with increased allocation of excitatory inputs onto inhibitory neurons and diminished convergence onto excitatory neurons (**Figs. 6J, 7G**), limiting the risk of over-excitation. This arrangement resembles homeostatic E/I plasticity in which excitatory synapses onto PV interneurons are up-scaled while those onto excitatory neurons are down-scaled to prevent runaway excitation ^62,63^. In AD-Impaired rats, HPC excitatory connectivity instead becomes excessive. Excitatory neurons receive highly convergent excitatory input (**Figs. 6H, 6I, 7G**), promoting hyperexcitation that is not adequately compensated by inhibitory control. In aging impairment, hippocampal excitatory failure manifests as hypoexcitable (**Fig. 4C**), with more pronounced synaptic loss than in AD impairment (**Fig. 6H**). These region-specific patterns suggest that maladaptive hyperconnectivity drives HPC hyperexcitability in AD dementia, whereas functional synaptic loss drives hypoactivity in aging-related impairment.

### Inhibitory circuits – S1FL

Differences in inhibitory circuitry further separate AD resilience from healthy aging and impairment. In S1FL of AD-Resilient rats, inhibitory divergence onto excitatory neurons is reduced relative to healthy aging (**Fig. 7E**), indicating that inhibitory control is applied more selectively. Under this configuration, the recruitment reliability of cortical excitatory neurons depends more strongly on excitatory synaptic convergence than in healthy aging (**Fig. 7K**), suggesting a compensatory mechanism that preserves stable cortical output despite AD pathology and altered interneuron function. In AD-Impaired cortex, we infer that inhibitory recruitment fails to counteract heightened excitatory drive. Although more neurons may be recruited overall, the lack of stable excitatory ensembles and the upregulated excitatory connectivity indicate ineffective inhibitory regulation.

### Inhibitory circuits – HPC

HPC Inhibitory circuits diverge sharply across healthy aging, AD resilience, AD dementia, and aging impairment. In healthy aging, inhibitory neurons form numerous synaptic connections with substantial divergence onto excitatory neurons (**Figs. 6I, 7E, 7I**), providing precise regulation of excitatory activity. In AD resilience, inhibitory control is reorganized rather than simply lost (**Fig. 7H**). Increased excitatory drive onto inhibitory neurons coupled with fewer excitatory inputs per interneuron (**Figs. 6J, 7H**) likely biases the network toward frequent but efficient interneuron recruitment, yielding stable inhibitory population representations as in healthy aging (**Fig. 5**). In ‘AD Resilient’ rats, HPC inhibitory neurons with more presynaptic excitatory partners are more reliably engaged than in healthy aging (**Fig. 7L**), and reduced inhibitory divergence onto excitatory neurons tuned to reduced excitatory convergence (**Figs. 7I, 7G**) may establish a new E/I balance that stabilizes network output.

In AD dementia, this rebalancing may fail. Both somatostatin (SST) and parvalbumin (PV) interneurons are vulnerable to AD pathologies ^64–66^, with SST interneurons affected earlier^67^. PV interneurons in TgF344-AD rats show early upregulation and increased dendritic complexity ^67^, and we observe a larger pool of inhibitory neurons recruited by repeated stimulation in ‘AD Impaired’ than in ‘AD Resilient’ rats (**Fig. 4C**) despite similar total GAD67 □NeuN □ counts (**Supplementary Fig. 3B**), suggesting hyperexcitable, over-recruited PV cells attempting to compensate for SST loss. Furthermore, loss of SST-mediated dendritic gating promotes excessive excitatory connectivity ^66,68^, weaker dependence of inhibitory engagement on excitatory convergence (**Fig. 7L**), more variable inhibitory subsets across trials (**Fig. 5**), and increased inhibitory divergence onto other inhibitory neurons (**Fig. 7J**), leading to disinhibition.

Postsynaptic inhibitory neurons also receive more excitatory inputs (**Fig. 7G**), culminating in overwhelming hyperexcitation in AD-Impaired rats.

In aging impairment, HPC inhibitory failure takes a different form: a hypoactive network with greater synaptic loss (**Fig. 6H**), reduced excitatory input onto inhibitory neurons (**Fig. 6I**), and diminished recruitment of inhibitory cells (**Fig. 4C**). Loss of inhibitory control in this context may exacerbate excitatory overdrive when stimuli are present and likely contributes to unstable population representations.

In summary, this study established a high-resolution electrophysiological phenotype associated with cognitive resilience in Alzheimer’s pathology. Our findings suggest that stabilizing neuronal representations may represent promising therapeutic strategies for preserving cognition in aging and neurodegenerative diseases.

## Supporting information

Statistical Tables

## Acknowledgments

We express our gratitude to Jessica Ribeiro and Matthew Mandrozos for their kind assistance in immunohistochemical experiments and imaging. This work has been funded by Canadian Institutes of Health Research (CIHR PJT191806) and National Institutes of Health (NIH R01AG084681). Dr. Keying Chen is supported by the Canadian Consortium on Neurodegeneration in Aging (CCNA) postdoctoral fellowship award.

## Data and code availability

All data and code will be made available on the Borealis platform at the time of publication.

## Methods

### Experimental model: TgF344-AD rats and non-transgenic littermates

This study is a cross-sectional study including a total of 53 rats at 13 months of age: 28 males and 25 females (**Fig. 1A**). Non-transgenic (nTg) and TgF344-AD rats were bred in-house, and maintained under a 12-h light–dark schedule with ad libitum access to chow and water. The TgF344-AD line models Alzheimer’s disease pathology through overexpression of human amyloid precursor protein with the Swedish mutation (APP KM670/671NL) and presenilin 1 (PS1) with exon 9 excised. Rats from multiple litters were assigned to groups according to genotype and behavioral performances. Experimental protocols were approved by the Animal Care Committee of the Sunnybrook Health Sciences Center, which adheres to the guidelines and policies of the Canadian Council on Animal Care and meets the Provincial Statute of Ontario, Animals for Research Act, and Federal Health of Animals Act.

### Barnes maze

To evaluate learning, spatial memory, and executive function in both transgenic (Tg) and non-transgenic (nTg) rats at the symptomatic stage (13 months) and young adulthood (4 months), prior to amyloidogenesis, Barnes maze test (Maze Engineers, Boston, MA) was conducted. All rats were behaviorally naïve prior to behavioral assessment. Barnes maze (Maze Engineers, Boston, MA) testing was conducted in a behavioral suite equipped with spatial cues, using an aversive light in all trials except during training (**Fig. 1B**). Video recordings were collected and analyzed with EthoVision XT (Noldus, Wageningen, Netherlands; version 11.5). After initial training to the escape location, rats underwent learning trials over 3 consecutive days (2 trials per day). Spatial memory was evaluated in a single probe trial conducted 3 days later. Reversal learning to assess executive function and cognitive flexibility began the next day and consisted of 5 days of testing (2 trials per day). During reversal learning, the escape hole was relocated without additional training. Task performance was quantified by measuring the latency to reach the escape hole.

### *In vivo* electrophysiological recordings Acquisition

Rats were initially anesthetized with 5% isoflurane for induction and then transferred to a stereotaxic frame. Anesthesia was maintained with 2–2.5% isoflurane, and physiological readouts including heart rate, respiration rate, and pO □ were continuously monitored (MouseOx Plus, Starr Life Sciences Corp., Oakmont, PA, USA). Core body temperature was maintained at 37 ± 1°C using a rectal probe and heat pad connected to a feedback system (TC-1000, CWE Inc., Ardmore, PA, USA). A 2-mm-square craniotomy was performed over the right somatosensory forepaw area (S1FL), centered at AP −0.9mm and ML +4.5mm. A second 2-mm-square craniotomy centered at AP +2.8 mm and ML +2.0 mm was performed for hippocampal recording. Both procedures were performed with care to avoid major surface vessels. Then the rat was transferred to a Faraday cage, and a Neuropixels 1.0 silicon probe was inserted at 3.5 µm/s with a four-axis manipulator (uMp-4, Sensapex) along a trajectory passing through the cortical S1FL region and hippocampus. Forepaw stimulation was delivered via subcutaneous electrodes placed in the left forepaw, as previously described ^69^. Each stimulation trial consisted of a 60-s train of 3 Hz pulses (0.3 ms duration, 10 mA intensity; 187 pulses total, BioPac MP150 System), followed by 300 s of no stimulation. This sequence was repeated 10 times per animal. Neuropixels 1.0 probes were used to acquire extracellular recordings both during stimulation and during a 10-min spontaneous baseline period prior to stimulation onset (**Fig. 1C**). The electrophysiological signals were recorded with a PXIe acquisition module (IMEC) and visualized and stored using Open Ephys software.

### Electrophysiology data analysis

Laminar alignment of the Neuropixels 1.0 probe was performed for every animal using current source density (CSD) analysis. A 2nd-order Butterworth filter at 0.4-300 Hz was applied to obtain the local field potential (LFP) data stream. LFP signals were smoothed by running 1-dimensional line fit. Then CSD was constructed by computing the second spatial derivative of evoked (stimulus-locked) LFP averaged from each 4 electrode sites 80 µm spacing along the microelectrode shank. CSD was averaged across 10 trials of forepaw stimulation, and the locations of the current sink-source couples within 100 ms were used to determine S1FL and hippocampus regions.

### Local field potential power analysis

LFP spectral power was calculated as described previously. For each condition (baseline and forepaw stimulation), power spectra were computed in MATLAB using the ‘spectrogram’ function with 2-s windows, a Hann taper, and 50% overlap to minimize spectral leakage. Power values were converted to decibels (dB) by logarithmic transformation. The normalized changes in power were calculated as the difference in mean power between stimulation and baseline across six frequency bands: delta (1–4 Hz), theta (4–8 Hz), alpha (8–12 Hz), beta (12–30 Hz), low gamma (30–60 Hz), and high gamma (60–90 Hz).

### Single-unit sorting and putative subtype classification

Raw spike stream data (0.3–10 kHz) were processed using SpikeInterface. Preprocessing included phase-shift correction, bandpass filtering (0.5–6 kHz), and common average referencing (CAR) before spike sorting with Kilosort 4 ^70^. Single units (SUs) were manually curated in Phy based on waveform shapes, auto-correlograms, and peri-stimulus time histograms (PSTHs). The three-dimensional spike localization model was implemented following previously described methods, using an open-source analysis package ^71^. Specifically, by applying a point-source model of voltage decreases with distance from the neuron, the spatial origin of SU clusters was estimated based on the peak-to-peak amplitudes of waveforms recorded across multiple sites on the Neuropixels probe^71^. SU putative subtype classification followed established waveform criteria ^72^: the waveform width was defined as the trough-to-peak (TP) latency, measured as the interval between the minimum (valley) and maximum (peak) of each averaged waveform. The bimodal distribution of TP latencies was modeled as a mixture of two Gaussian distributions (μ_1_ = 0.31 ms, σ □ = 0.05 ms; μ_2_ = 0.48 ms, σ_2_ = 0.07 ms), and the intersection point (0.38 ms) was used as the classification threshold. SUs with TP latency > 0.38 ms were classified as putative excitatory neurons, and those with TP latency ≤ 0.38 ms as putative inhibitory neurons. Signal-to-noise ratios (SNR) were calculated by dividing the peak-to-peak amplitude of each single unit by the noise.

### Changes in Inter-Spike Interval

To assess changes in spike timing dynamics, we computed Inter-Spike Interval (ISI) histograms during both baseline and sensory stimulation trials. For each neuron, the ISI of spike n was plotted against the ISI of spike n + 1 to visualize sequential firing variability. ISI histograms were generated separately for the forepaw stimulation and baseline conditions, and their difference (stimulation – baseline) was calculated to obtain the □ ISI histogram. All ISI difference histograms were then averaged across responsive neuronal subtypes and generate a mean □ ISI histogram representative of each group. Statistical comparisons of mean histograms between groups were performed using a permutation test (12,500 permutations).

### Spiking firing irregularity measurements

Spiking irregularity was quantified by analyzing the sequence of inter-spike intervals (ISIs) for each unit. An enhanced local variation measure, Local Variation with Refractoriness correction (LvR), was used to assess firing regularity within single trials as previously described ^51^. In contrast to conventional measures such as the coefficient of variation (CV), LvR accounts for fluctuations in firing rate across the spike train and corrects for the influence of the refractory period following each spike. LvR is defined as:

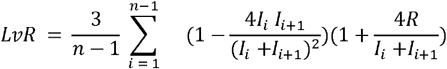

where *I*_*i*_ denotes the i-th ISI, *n* is the total number of ISIs, and *R* is the refractoriness constant and was set to 5ms based on the empirical calibration reported by Shinomoto et al. (2003) ^51^. LvR values were computed from spike sequences containing at least five spikes to ensure statistical reliability; sequences not meeting this criterion were excluded from analysis.

### Neuronal representations measurements

To assess the stability of neuronal representations across repeated sensory stimuli, we defined the Neuronal Engagement Index (NEI). Each trial *t* contains *P* = 187 stimulation pulses, followed by 300 s of resting. Each animal performs *T* = 10 trials. Let *r*_*i*,*p*,*t*_ ∈ {0,1} indicate whether neuron *i* exhibits a significant stimulus-evoked change in firing rates on pulse *p* of trial *t* compared to its pre-stimulation baseline (permutation test, p<0.05). Let *A*_*i*,*t*_ ∈ {0,1} indicate if neuron *i* is activated in trial *t*. Let 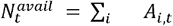 be the number of neurons that could be activated within that trial. Therefore, for a given neuron *i* in trial *t*, the fraction of the 187 stimulus pulses that neuron *i* elicits significant responses is:

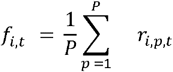

We also noted that for a given stimulus pulse *p* of trial *t*, the fraction of available neurons significantly engaged by that stimulus pulse is:

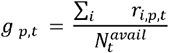

We observed that averaging *f*_*i*,*t*_ across available neurons and averaging *g* _*p*,*t*_ across stimulus pulses yield the same trial-level quantity, which we define as the NEI:

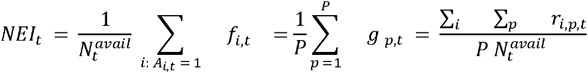

Thus, NEI of each trial simultaneously described the (1) proportion of neurons within a local network that were reliably activated by the stimulus and (2) reliability of a neuron’s response across repeated stimuli. The mean NEI was computed by averaging across trials and then across animals, separately for putative excitatory and putative inhibitory neuronal populations. NEI ranged from 0 to 1, where higher values indicated greater stability of neuronal representations. To determine the relationship between NEI and cognitive performance, we computed the Pearson’s correlation coefficient between NEI and each behavioral AUC metric, as well as with a weighted composite score (0.2·Learning t1-t6 AUC + 0.3·Reversal Learning t1-t6 AUC + 0.5·Reversal Learning t7-t10 AUC).

### Detection of monosynaptic functional connections

Cross-correlograms (CCGs) of spike trains were used to identify putative monosynaptic connections between neuron pairs. The CCG was computed as the histogram of time differences between spike occurrences in the reference and target spike trains within a window of −30 to +30 ms using 0.5-ms bins. Then the CCG histogram was convolved with a Gaussian kernel (σ = 7 ms) to estimate the expected spike probability and smooth random fluctuations. For each bin, the 99.9999th percentile of the cumulative Poisson distribution was calculated using the estimated average spike count from the convolved CCG. This serves as the statistical threshold for detecting significant deviations. A putative synaptic connection was identified when at least two consecutive bins within the 1.5–4 ms post-spike interval exceeded the upper Poisson threshold or fell below the corresponding lower Poisson threshold. Positive deflections indicated putative excitatory connections, and negative deflections indicated inhibitory connections. Waveform-based classification was further used to validate the CCG-identified presynaptic units. Based on these criteria, 10,398 cortical and 1,340 hippocampal connections were classified as putative excitatory monosynaptic pairs, while 1,231 cortical and 324 hippocampal connections were classified as putative inhibitory monosynaptic pairs. Connection strength represents the efficacy of spike transmission between each pair of cells and was defined as the standardized peak (or trough) height in the CCG. It was calculated as the absolute difference between the observed spike count in the peak (or trough) and the baseline mean, divided by the baseline standard deviation.

### Immunofluorescence and immunohistochemistry

Upon completion of electrophysiological recordings, rats were transcardially perfused under 5% isoflurane with phosphate-buffered saline (PBS; 0.1% heparin), followed by PBS containing 4% paraformaldehyde (PFA). Brains were extracted, post-fixed overnight in PBS-4% PFA, and cryoprotected in PBS-30% sucrose before microtome sectioning. For each brain, four 40-µm coronal sections containing both cortical and hippocampal regions were sampled. Imaging for all experiments was performed using a Zeiss Observer Z1 microscope (**Fig. 1D**).

### Tau Immunohistochemistry (PHF1)

Free-floating sections were washed in Tris-buffered saline (TBS) and incubated for 30 min at room temperature in 3% hydrogen peroxide with 0.25% Triton X-100 in TBS to quench endogenous peroxidase activity. Sections were then blocked in 5% skim milk in TBS for 1h and incubated overnight at 4 °C with mouse anti-PHF1 (1:1000; courtesy of the late Dr. Peter Davies, The Feinstein Institute for Medical Research). The following day, sections were incubated with Biotin-XX goat anti-mouse IgG1 (1:200; ThermoFisher A10519, Lot #2168874), followed by avidin-biotin complex (ABC) and visualized using 3,3′-diaminobenzidine (DAB; Vector Labs) without nickel enhancement. Sections were dehydrated in graded ethanols, cleared in xylene, and mounted with Cytoseal XYL. Quantification of tau was performed in ImageJ Fiji by manual counting. The density of PHF1-positive inclusions was calculated as the number of tau pre-tangles per mm^2^ in hippocampal and cortical regions separately.

### β*-Amyloid Immunofluorescence*

Sections underwent antigen retrieval in 10 mM sodium citrate buffer (pH 6.0, 90 °C, 30 min), were blocked in PBS containing 5% goat serum, 0.25% Triton X-100, and 0.25% BSA, and incubated overnight at 4 °C with rabbit β-amyloid (D54D2) XP antibody (1:300; Cell Signaling #8243). Sections were then incubated for 2 h at room temperature with goat anti-rabbit IgG Alexa 488 (1:250; ThermoFisher A11008), mounted with PVA-DABCO, and coverslipped. β-Amyloid coverage was quantified in ImageJ Fiji by thresholding and binarizing images to determine staining density.

The percentage area occupied by β-amyloid was calculated for hippocampal and cortical regions separately.

### NeuN and GAD67 Immunofluorescence Co-staining

Sections were blocked in TBS containing 2% donkey serum, 0.5% Triton X-100, and 0.5% bovine serum albumin (BSA) for 1 h at room temperature, then incubated overnight at 4 °C with guinea pig anti-NeuN (1:500; Millipore ABN90) and mouse anti-GAD67 (1:250; Millipore MAB5406). The following day, sections were incubated with donkey anti-mouse IgG-HRP (1:500 Jackson Immunoresearch, #715-035-151) and goat anti-guinea pig IgG Alexa 594 (1:250 Thermo Fisher A-11076). Sections were mounted with PVA-DABCO mounting medium pre-warmed to 60 °C and coverslipped. Cell segmentation was performed in ilastik (all features enabled), and subsequent analysis was conducted in ImageJ Fiji. Densities of NeuN- and GAD67-positive cells were calculated as counts per mm^2^ within CA1, CA3, and cortical regions (particle size threshold = 0.00002 mm^2^ − ∞).

### Statistics

A linear mixed-effects (LME) model was applied to assess the effects of genotype and cognitive status on electrophysiological metrics. The model was implemented using MATLAB’s built-in *fitlme* function from the Statistics and Machine Learning Toolbox. The LME examined the relationship between the dependent variable (e.g., neuronal engagement index, NEI) and two fixed factors (genotype and cognitive status) while accounting for random inter-animal variability. This model has the form: NEI = β0 + β1 * Genotype + β1 * Cognitive Status + (1∣Rat ID) + □, where β0 is the intercept, β1 and β2 represented the fixed effect coefficients that represents the average change in response variable (NEI) associated with genotype and cognitive status. (1∣Rat ID) denotes the random intercept accounting for variability across individual rats. ε is the residual error term. For immunohistochemical analyses, a Welch’s test was used to determine significant differences in D54D2 area coverage between Tg cognitively resilient and impaired rats. Additionally, a two-way ANOVA (p < 0.05) was used to assess the effects of genotype and cognitive status on densities of PHF1, NeuN, and GAD67. Tukey’s post hoc test (Least Significant Difference correction) was applied to determine pairwise group differences. Pearson’s correlation coefficient (r) was used to assess the association between neuronal convergence and the neuronal engagement index (NEI), and the corresponding p-value was reported to indicate statistical significance. ANCOVA analyses were further conducted to test whether these relationships differed across the four groups stratified by genotype and cognitive status. All p-values are reported in Supplementary Tables.

---

**Supplementary Figure 1:**
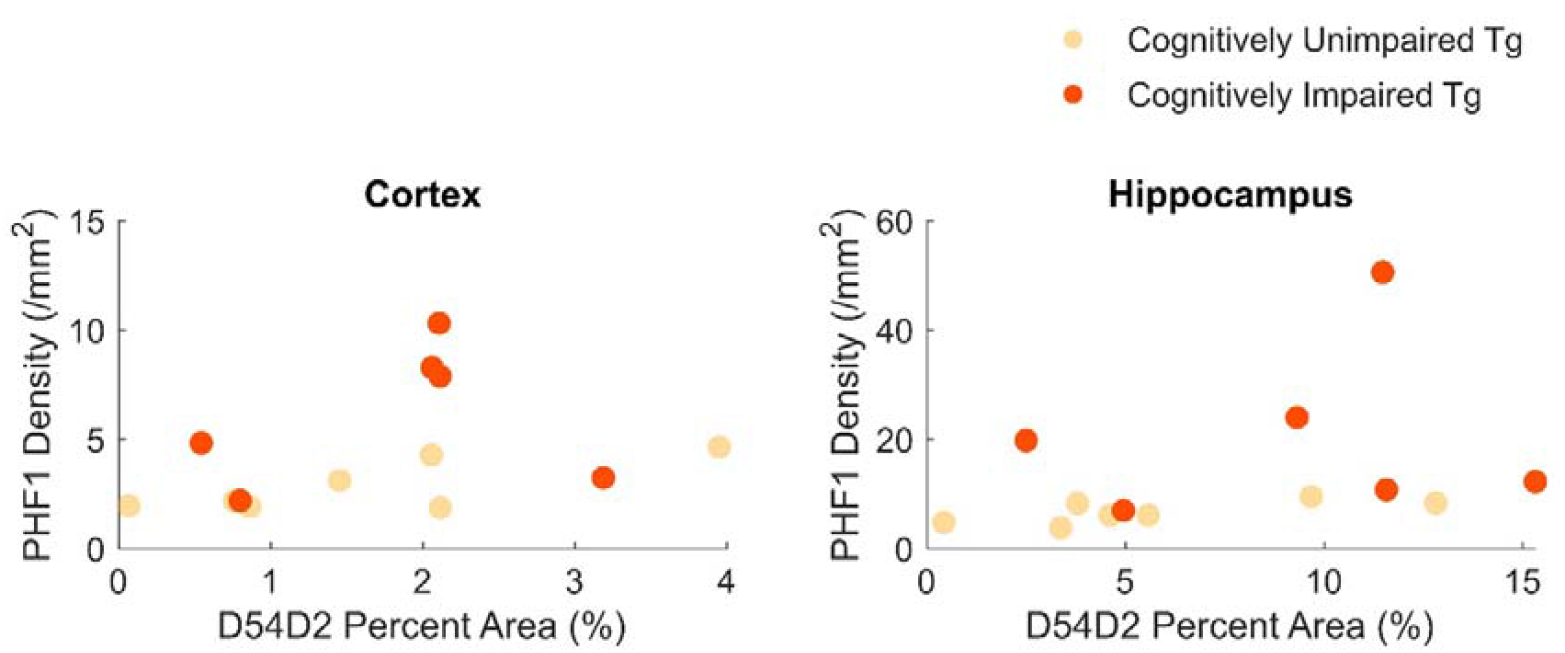
Pathological hallmarks are not sufficient to determine cognitive status of Fischer 344 TgAD rats. Scatter plots for Fischer 344 TgAD rats show PHF1-positive inclusion density and D54D2 percent area coverage in the cortex and hippocampus. No clear separation was observed between cognitively unimpaired and impaired groups.

**Supplementary Figure 2:**
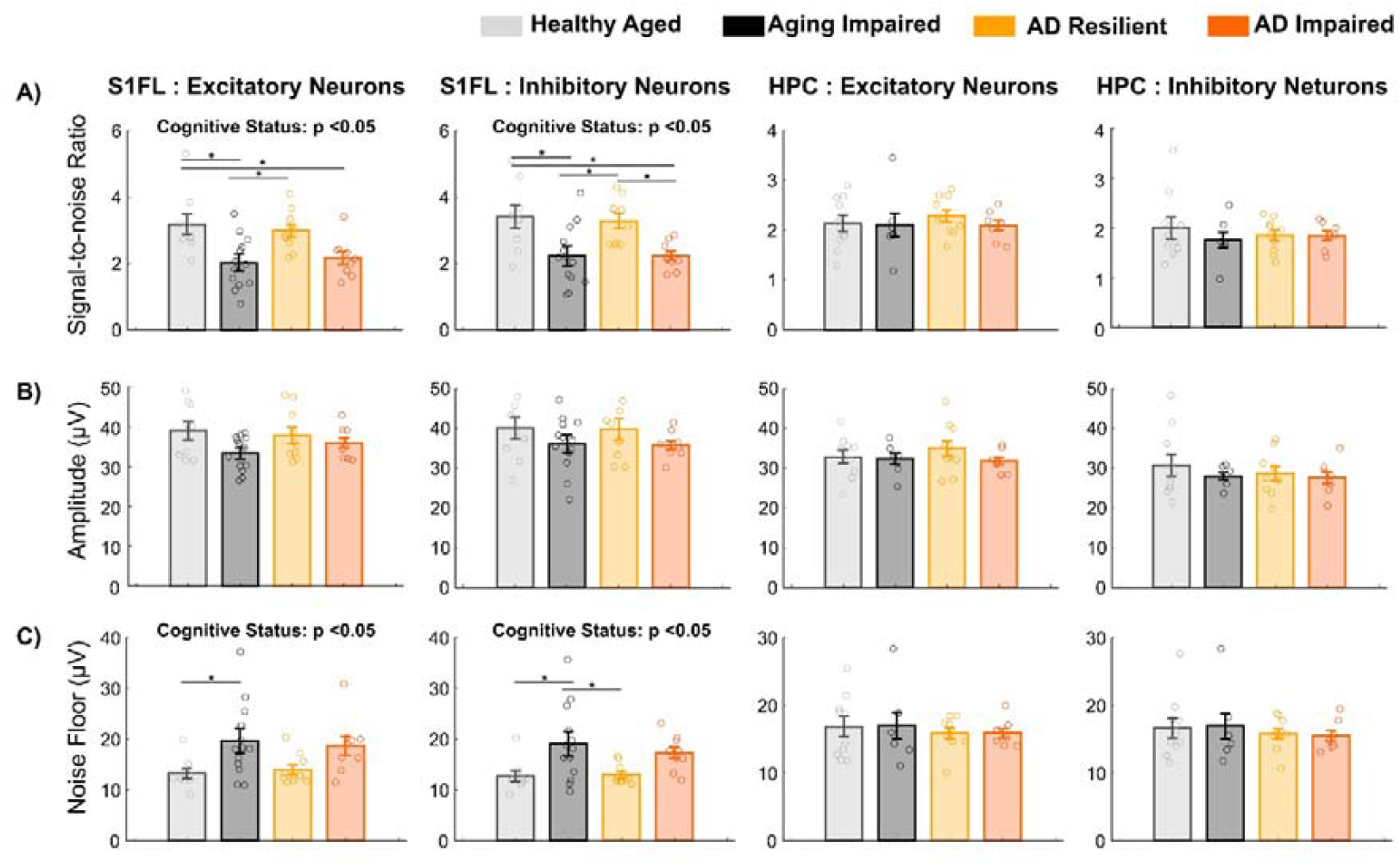
Ultra-High-density single-unit recordings exhibited high SNR in S1FL in cognitively resilient rats. **A)** S1FL excitatory and inhibitory neurons in ‘AD Resilient’ (E: 2.99 ± 0.59; I: 3.28 ± 0.68) and ‘Healthy Aged’ (E: 3.18 ± 0.93; I: 3.41 ± 1.02) rats had significantly higher SNR in than those in ‘Aging Impaired’ (E: 2.02 ± 0.78; I: 2.22 ± 0.90) and ‘AD Impaired’ (E: 2.17 ± 0.58; I: 2.23 ± 0.42) rats. **B)** peak-to-peak amplitudes were comparable for excitatory and inhibitory SU signals in S1FL and HPC. **C)** Noise levels in channels with detected SUs in S1FL were significantly lower in ‘AD Resilient’ (E: 14.01 ± 2.89; I: 13.04 ± 2.01) and ‘Healthy Aged’ (E: 13.27 ± 2.94; I: 12.77 ± 3.08) rats than in ‘Aging Impaired’ (E: 19.69 ± 7.33; I: 19.09 ± 7.44) and ‘AD Impaired’ (E: 18.71 ± 5.48; I: 17.38 ± 3.47) rats.

**Supplementary Figure 3:**
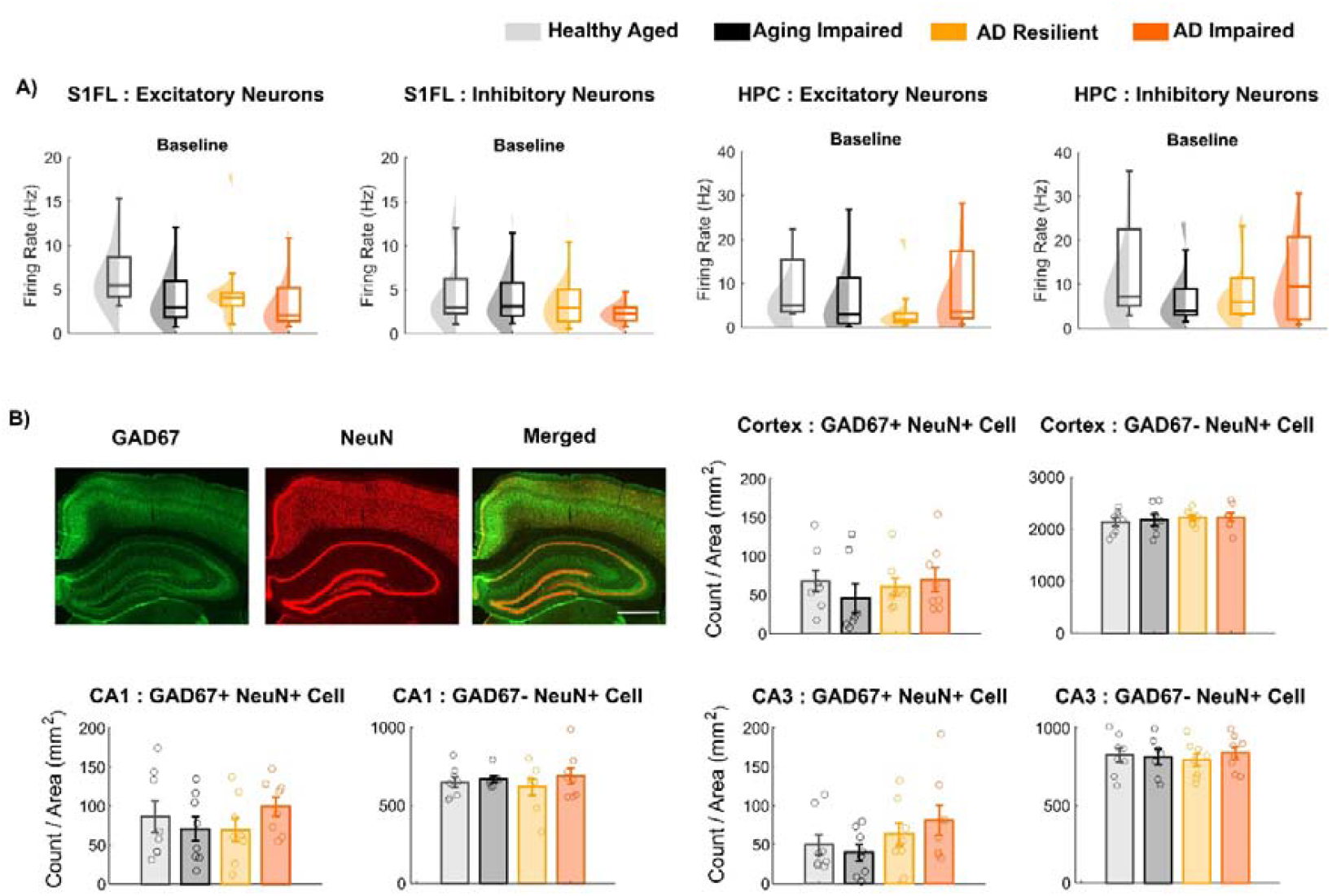
Alignment of recording sites and additional characterizations of Single Unit activities. **A)** The firing rates during pre-stimulation baseline were indistinguishable across the 4 groups for either putative excitatory or inhibitory neurons in S1FL and hippocampus. **B)** Immunostaining-based analysis showed that the densities of GAD67+NeuN+ (inhibitory) and GAD67−NeuN+ (excitatory) neurons in cortex, CA1, and CA3 were comparable across genotypes and cognitive groups. Scale bar = 1 mm.

**Supplementary Figure 4:**
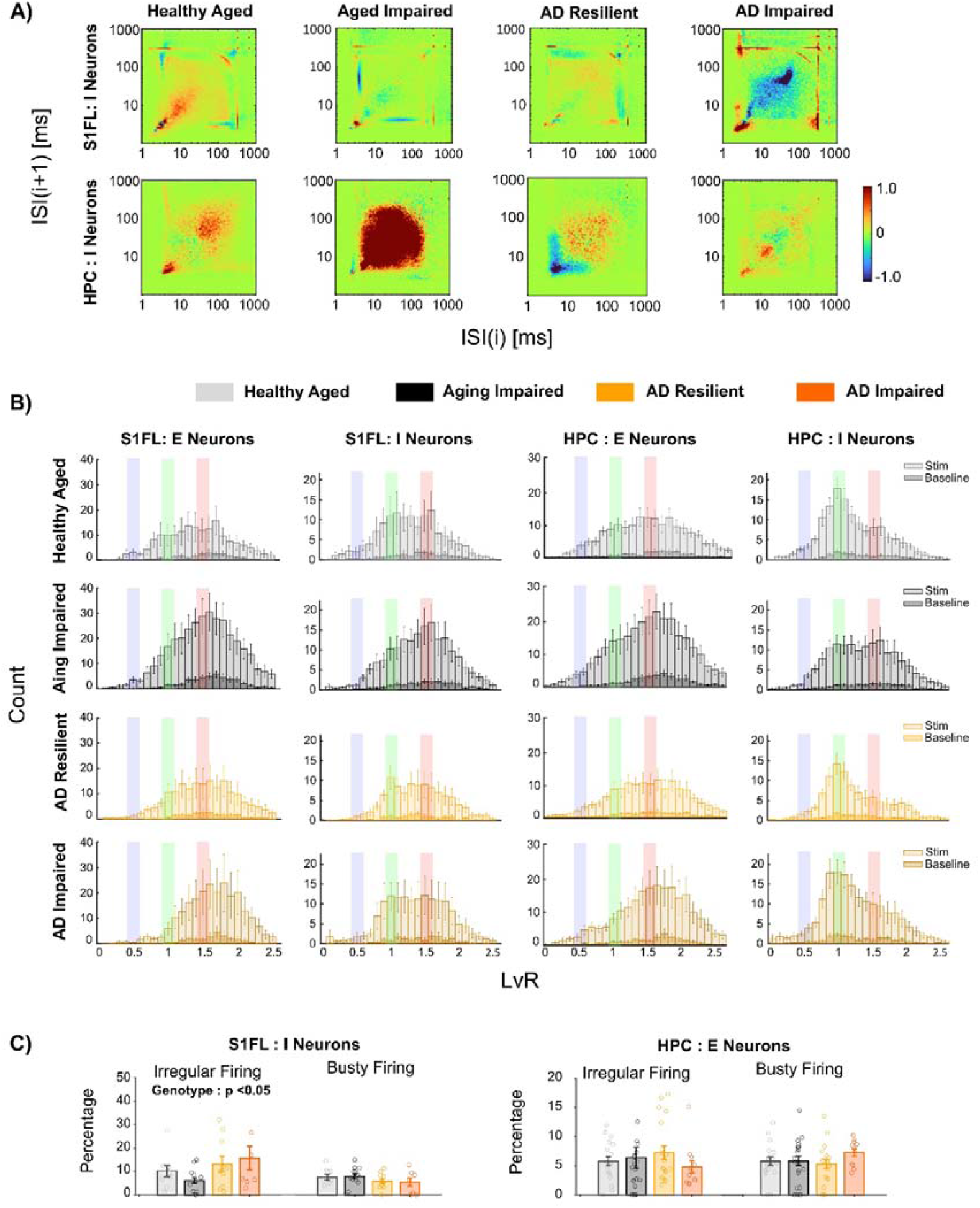
Patterns of spike dynamics during network activation. **A)** In S1FL, ‘AD Impaired’ rats exhibited a selective reduction in ΔISI spike probability at ~40 ms. In the hippocampus, ΔISI histograms indicated increased burstiness during network activation in ‘Aging Impaired’ rats. **B)** Histograms of LvR values during forepaw stimulation and pre-stimulation baseline. Colored overlays represent the range of +0.05 around 0.5 (blue, regular firing), 1.0 (green, random firing), and 1.5 (red, bursty firing). **C)** In S1FL, the percentage of irregular-firing inhibitory neurons (LvR ≈ 1) showed a significant main effect of genotype, with Tg rats (AD Resilient: 13.21 ± 10.78%, AD Impaired: 15.65±15.02%) exhibiting a significantly higher percentage than nTg rats (Healthy Aged: 10.17 ± 7.47%, Aging Impaired: 6.07 ± 4.69%).

**Supplementary Figure 5:**
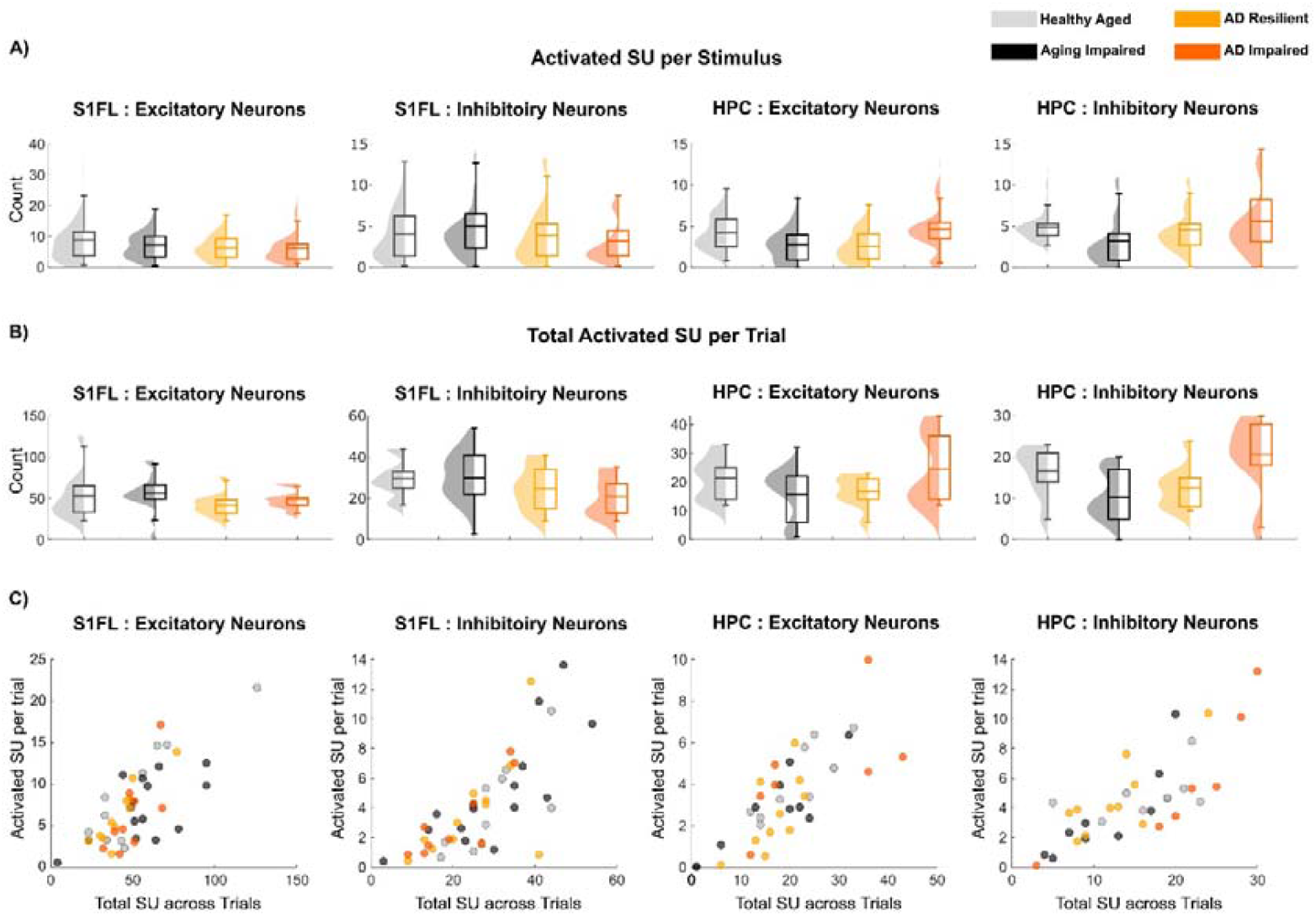
Neuronal representation variability. For both putative excitatory and inhibitory neurons in S1FL and hippocampus, **A)** the number of activated single units per stimulus and **B)** The total neurons activated per stimulation trial (with 187 stimuli per trial) were statistically comparable among the four groups. **C)** The mean number of activated SUs per stimulus against the total SU across all trials of each animal.

**Supplementary Figure 6:**
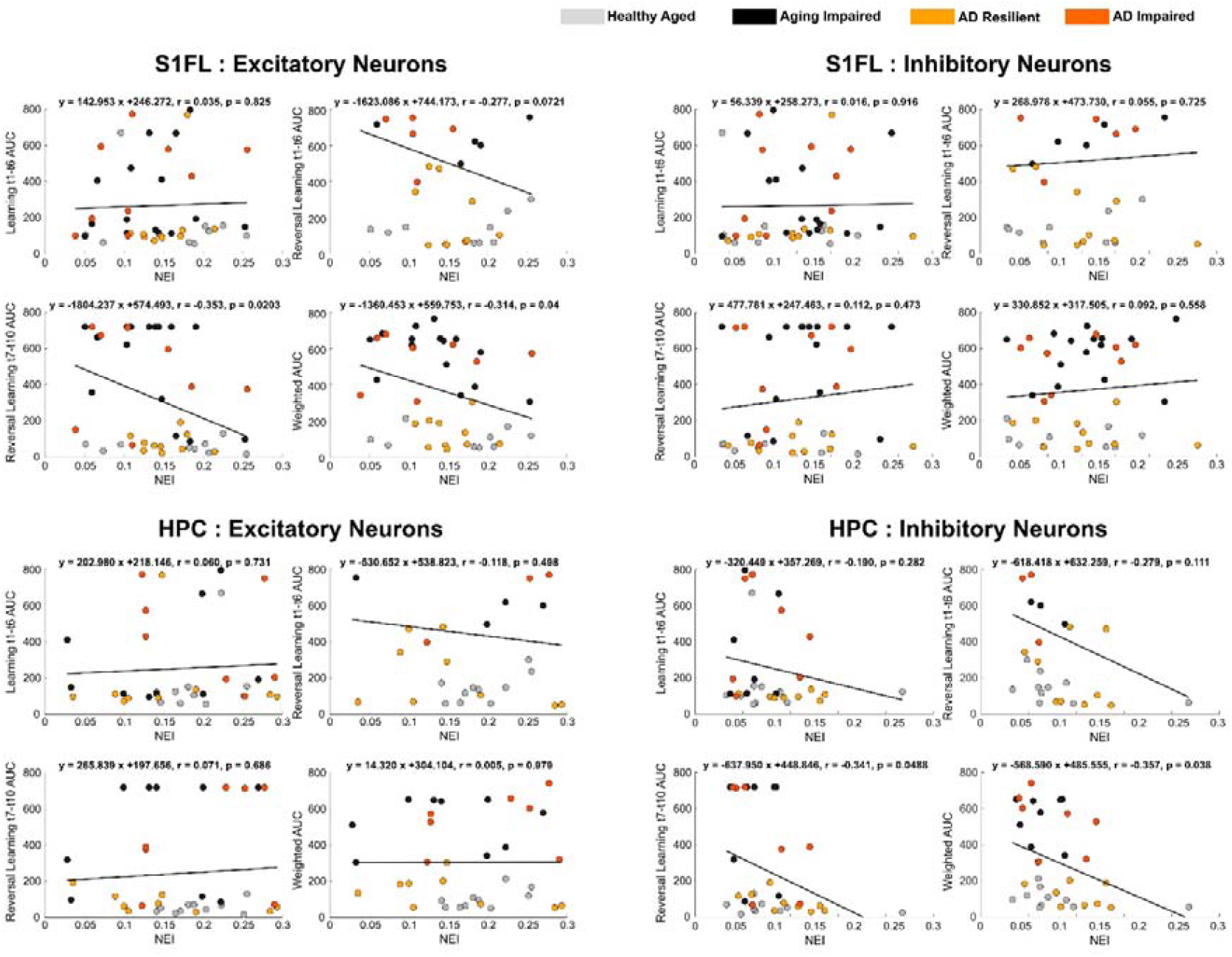
Correlation between NEI and Cognitive Performances. Pearson correlation coefficients and regression slopes are shown for the relationships between NEI and the latency AUC metrics (Learning t1–t6, Reversal Learning t1–t6, Reversal Learning t7–t10), as well as a weighted composite score.

**Supplementary Figure 7.**
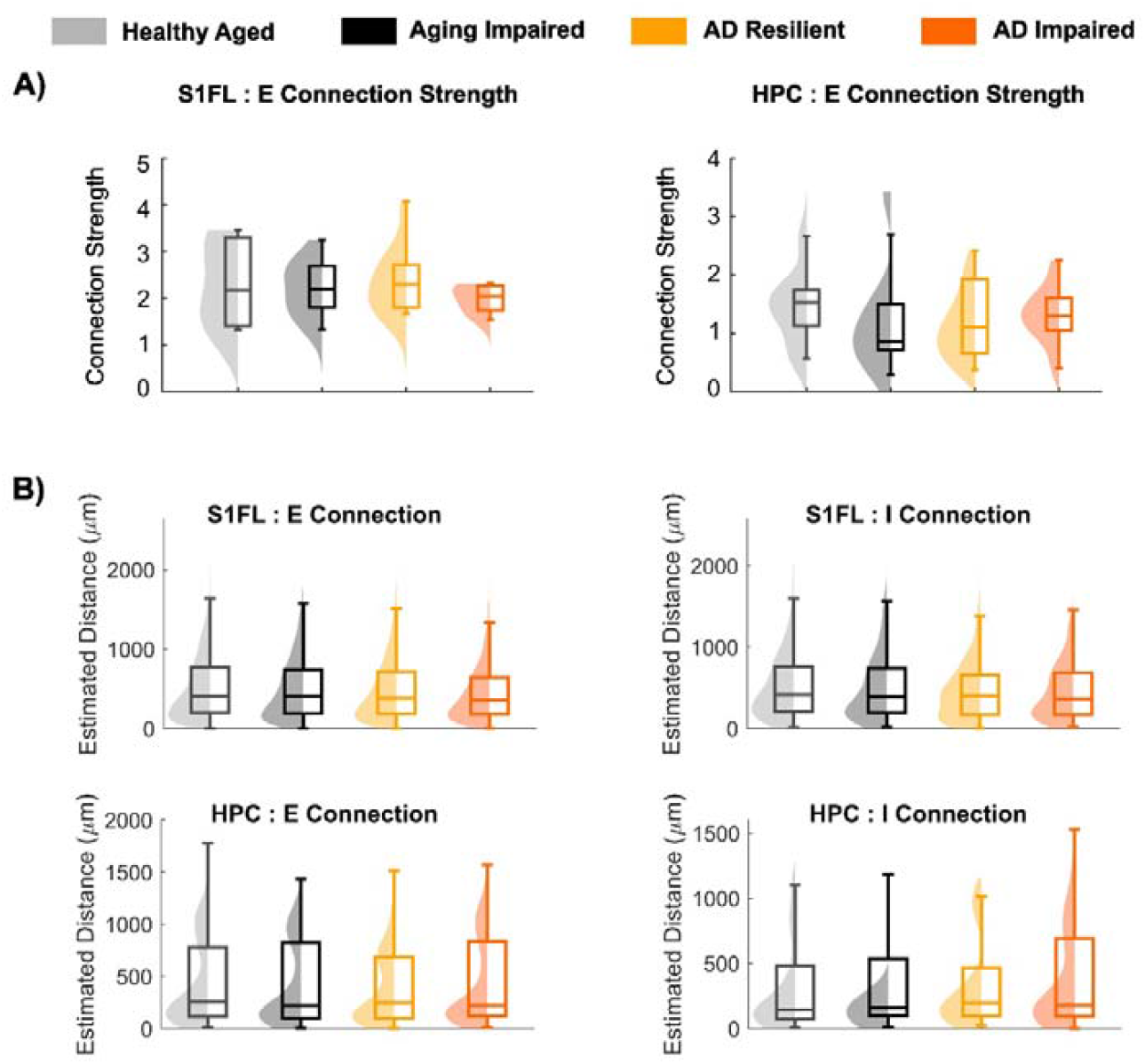
Characterization of CCG-based synaptic connections. **A)** Strength of CCG-based excitatory synaptic connections were comparable across genotypes and cognitive-status groups in both S1FL and hippocampus. **B)** Intersomatic distance between pre- and postsynaptic neurons were estimated from three-dimensional spatial coordinates reconstructed using the high-density recording sites of the Neuropixels probe. No significant differences in the distances between neurons for CCG-based excitatory or inhibitory connection pairs were observed across genotypes or cognitive status in either S1FL or the HPC.

**Supplementary Figure 8:**
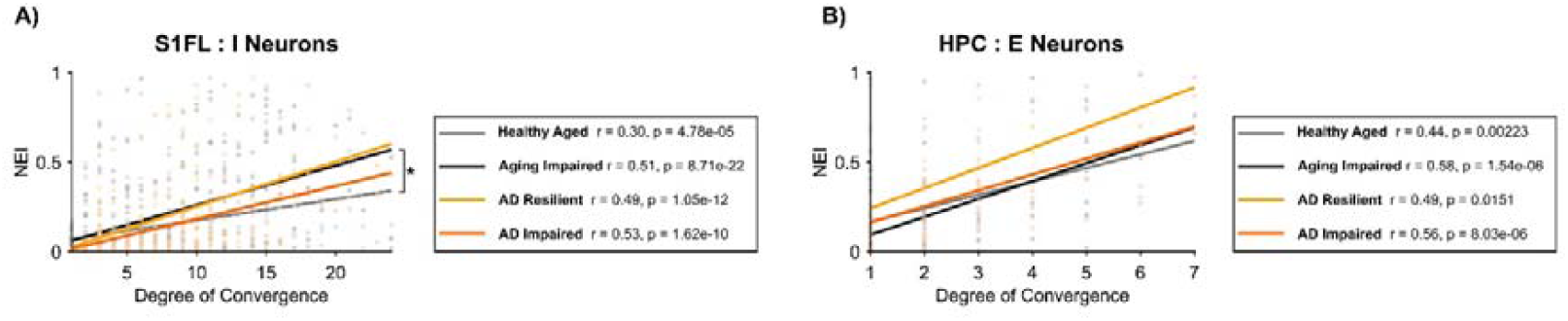
Convergence–NEI relationships. A) In S1FL, the linear regression of the degree of convergence onto putative inhibitory neurons vs. their NEI showed a significantly steeper slope in ‘Aging Impaired’ rats vs. in ‘Healthy Aged” rats. **B)** In the hippocampus, the coefficients of Pearson’s correlations between the degree of excitatory inputs convergent onto HPC excitatory neurons and their NEI were comparable across genotype and cognitive-status groups.

