## Supplementary material for "Stable neuronal representations to repeated stimulation underlie cognitive resilience in Alzheimer’s disease pathology": Statistical Tables

| Barnes Maze Latency | | Learning | Reversal Learning |
| --- | --- | --- | --- |
| Model | Variable | p-value | p-value |
| Linear Mixed Model | Age | 0.0976 | 0.615 |
|  | Genotype | 0.199 | 0.200 |
|  | Session Num | 0.430 | ***0.000289*** |
|  | Age:Genotype | 0.267 | 0.318 |
|  | Age: Session Num | 0.727 | ***0.0421*** |
|  | Genotype: Session Num | 0.277 | 0.182 |
|  | Age:Genotype: Session Num | 0.119 | 0.316 |

**Supplementary Table 1.** Statistical analysis of escape latencies on Barnes Maze Learning and Reversal Learning phases; significance *p*<0.05; 4 mo nTg: n=12; 4 mo Tg: n=12; 13 mo nTg: n = 26; 13 mo Tg: n=27. Significant p-values are bolded and italicized.

| Area Coverage of D54D2 | | Cortex | HPC |
| --- | --- | --- | --- |
| Model | Variable | p-value | |
| Welch’s t-test | Cognitive Status | 0.770 | 0.459 |

**Supplementary Table 2.** Statistical analysis of D54D2 area coverage; significance *p*<0.05; ‘AD Resilient’ rats: n=7; ‘AD Impaired’ rats: n =8.

| Model | PHF1 Density: **Cortex** | | | | | | | | |
| --- | --- | --- | --- | --- | --- | --- | --- | --- | --- |
|  | Main effects | p-value | Group-wise | ‘Healthy Aged’ | | ‘Aging Impaired’ | ‘AD Resilient’ | | ‘AD Impaired |
| Linear Mixed Model + Tukey post hoc with LSD correction | Genotype | ***8.88e-4*** | ‘Healthy Aged’ | x | | 0.994 | 0.648 | | ***0.000317*** |
|  |  |  | ‘Aging Impaired’ | x | | x | 0.525 | | ***0.000254*** |
|  | Cognitive Status | 0.0508 | ‘AD Resilient’ | x | | x | x | | ***0.00809*** |
|  |  |  | ‘AD Impaired | x | | x | x | | x |
|  | PHF1 Density: **HPC** | | | | | | | | |
|  | Main effects | p-value | Group-wise | ‘Healthy Aged’ | ‘Aging Impaired’ | | ‘AD Resilient’ | ‘AD Impaired | |
|  | Genotype | ***7.45e-4*** | ‘Healthy Aged’ | x | 0.999 | | 0.669 | ***0.000287*** | |
|  |  |  | ‘Aging Impaired’ | x | x | | 0.627 | ***0.000325*** | |
|  | Cognitive Status | ***0.0376*** | ‘AD Resilient’ | x | x | | x | ***0.00679*** | |
|  |  |  | ‘AD Impaired | x | x | | x | x | |

**Supplementary Table 3.** Statistical analysis of PHF1 density; significance *p*<0.05; ‘Healthy Aged’: n=8; ‘Aging Impaired’: n=8; ‘AD Resilient’ rats: n=7; ‘AD Impaired’ rats: n =8. Significant p-values are bolded and italicized.

| Model | | Evoked LFP Power | | | | | | |
| --- | --- | --- | --- | --- | --- | --- | --- | --- |
|  |  | Main effects | p-values | | | | | |
|  |  |  | S1FL: Delta | S1FL: Alpha | HPC: Alpha | HPC: Beta | HPC: Low Gamma | HPC: High Gamma |
| Linear Mixed Model | | Genotype | 0.172 | 0.328 | 0.432 | 0.622 | 0.492 | 0.0653 |
|  |  | Cognitive Status | 0.332 | 0.224 | 0.436 | 0.624 | 0.474 | 0.452 |
| Model | Evoked LFP Power: S1FL Theta | | | | | | | |
|  | Main effects | | p-value | Group-wise | ‘Healthy Aged’ | ‘Aging Impaired’ | ‘AD Resilient’ | ‘AD Impaired |
| Linear Mixed Model + Tukey post hoc with LSD correction | Genotype | | 0.0576 | ‘Healthy Aged’ | x | ***0.0449*** | 0.911 | 0.975 |
|  |  |  |  | ‘Aging Impaired’ | x | x | ***0.00222*** | 0.116 |
|  | Cognitive Status | | ***0.00871*** | ‘AD Resilient’ | x | x | x | 0.672 |
|  |  |  |  | ‘AD Impaired | x | x | x | x |
|  | Evoked LFP Power: S1FL Beta | | | | | | | |
|  | Main effects | | p-value | Group-wise | ‘Healthy Aged’ | ‘Aging Impaired’ | ‘AD Resilient’ | ‘AD Impaired |
|  | Genotype | | 0.179 | ‘Healthy Aged’ | x | ***5.49e-5*** | ***3.38e-6*** | ***3.93e-7*** |
|  |  |  |  | ‘Aging Impaired’ | x | x | 0.213 | 0.534 |
|  | Cognitive Status | | ***1.25e-12*** | ‘AD Resilient’ | x | x | x | 0.956 |
|  |  |  |  | ‘AD Impaired | x | x | x | x |
|  | Evoked LFP Power: S1FL Low Gamma | | | | | | | |
|  | Main effects | | p-value | Group-wise | ‘Healthy Aged’ | ‘Aging Impaired’ | ‘AD Resilient’ | ‘AD Impaired |
|  | Genotype | | 0.161 | ‘Healthy Aged’ | x | ***1.81e-6*** | ***2.37e-6*** | ***1.20e-9*** |
|  |  |  |  | ‘Aging Impaired’ | x | x | ***0.0104*** | 0.499 |
|  | Cognitive Status | | ***3.15e-16*** | ‘AD Resilient’ | x | x | x | 0.428 |
|  |  |  |  | ‘AD Impaired | x | x | x | x |
|  | Evoked LFP Power: S1FL High Gamma | | | | | | | |
|  | Main effects | | p-value | Group-wise | ‘Healthy Aged’ | ‘Aging Impaired’ | ‘AD Resilient’ | ‘AD Impaired |
|  | Genotype | | 0.502 | ‘Healthy Aged’ | x | ***3.59e-10*** | ***0.0360*** | ***1.48e-4*** |
|  |  |  |  | ‘Aging Impaired’ | x | x | ***6.99e-4*** | 0.197 |
|  | Cognitive Status | | ***6.43e-11*** | ‘AD Resilient’ | x | x | x | 0.351 |
|  |  |  |  | ‘AD Impaired | x | x | x | x |
|  | Evoked LFP Power: HPC Delta | | | | | | | |
|  | Main effects | | p-value | Group-wise | ‘Healthy Aged’ | ‘Aging Impaired’ | ‘AD Resilient’ | ‘AD Impaired |
|  | Genotype | | ***0.0139*** | ‘Healthy Aged’ | x | ***0.00144*** | 0.999 | 0.881 |
|  |  |  |  | ‘Aging Impaired’ | x | x | ***0.00221*** | 0.0659 |
|  | Cognitive Status | | ***2.53e-4*** | ‘AD Resilient’ | x | x | x | 0.898 |
|  |  |  |  | ‘AD Impaired | x | x | x | x |
|  | Evoked LFP Power: HPC Theta | | | | | | | |
|  | Main effects | | p-value | Group-wise | ‘Healthy Aged’ | ‘Aging Impaired’ | ‘AD Resilient’ | ‘AD Impaired |
|  | Genotype | | ***0.0169*** | ‘Healthy Aged’ | x | ***0.00275*** | 0.811 | 0.999 |
|  |  |  |  | ‘Aging Impaired’ | x | x | ***0.0334*** | 0.0788 |
|  | Cognitive Status | | ***0.00663*** | ‘AD Resilient’ | x | x | x | 0.885 |
|  |  |  |  | ‘AD Impaired | x | x | x | x |

**Supplementary Table 4.** Statistical analysis of evoked LFP power; significance *p*<0.05; 10 stimulation trials repetition in each animal subject; ‘Healthy Aged’: n=10; ‘Aging Impaired’: n=15; ‘AD Resilient’ rats: n=15; ‘AD Impaired’ rats: n =15. Significant p-values are bolded and italicized.

| Model | SU signal-to-noise (SNR) ratio: S1FL Excitatory neurons | | | | | | | |
| --- | --- | --- | --- | --- | --- | --- | --- | --- |
|  | Main effects | p-value | Group-wise | ‘Healthy Aged’ | | ‘Aging Impaired’ | ‘AD Resilient’ | ‘AD Impaired |
| Linear Mixed Model + Tukey post hoc with LSD correction | Genotype | 0.939 | ‘Healthy Aged’ | x | | ***0.00449*** | 0.945 | ***0.0300*** |
|  |  |  | ‘Aging Impaired’ | x | | x | ***0.0167*** | 0.964 |
|  | Cognitive Status | ***1.35e-4*** | ‘AD Resilient’ | x | | x | x | 0.0894 |
|  |  |  | ‘AD Impaired | x | | x | x | x |
|  | SU SNR: S1FL Inhibitory neurons | | | | | | | |
|  | Main effects | p-value | Group-wise | ‘Healthy Aged’ | | ‘Aging Impaired’ | ‘AD Resilient’ | ‘AD Impaired |
|  | Genotype | 0.823 | ‘Healthy Aged’ | x | | ***0.00779*** | 0.986 | ***0.0174*** |
|  |  |  | ‘Aging Impaired’ | x | | x | ***0.0160*** | 0.999 |
|  | Cognitive Status | ***7.91e-5*** | ‘AD Resilient’ | x | | x | x | ***0.0333*** |
|  |  |  | ‘AD Impaired | x | | x | x | x |
|  | SU SNR: HPC | | | | | | | |
| Model | Main effects | p-values | | | | | | |
|  |  | Excitatory neurons | | | Inhibitory neurons | | | |
| Linear Mixed Model | Genotype | 0.633 | | | 0.742 | | | |
|  | Cognitive Status | 0.776 | | | 0.573 | | | |

**Supplementary Table 5.** Statistical analysis of SU signal-to-noise (SNR) ratio; significance *p*<0.05; ‘Healthy Aged’: n=10; ‘Aging Impaired’: n=15; ‘AD Resilient’ rats: n=15; ‘AD Impaired’ rats: n =10. Significant p-values are bolded and italicized.

| SU peak-to-peak amplitude | | | | | |
| --- | --- | --- | --- | --- | --- |
| Model | Main effects | p-values | | | |
|  |  | S1FL: Excitatory neurons | S1FL: Inhibitory neurons | HPC: Excitatory neurons | HPC: Inhibitory neurons |
| Linear Mixed Model | Genotype | 0.693 | 0.896 | 0.641 | 0.732 |
|  | Cognitive Status | 0.0832 | 0.0742 | 0.922 | 0.292 |

**Supplementary Table 6.** Statistical analysis of SU peak-to-peak amplitude; significance *p*<0.05; ‘Healthy Aged’: n=10; ‘Aging Impaired’: n=15; ‘AD Resilient’ rats: n=15; ‘AD Impaired’ rats: n =10. Significant p-values are bolded and italicized.

| Model | Noise Floor: S1FL Excitatory neurons | | | | | | | |
| --- | --- | --- | --- | --- | --- | --- | --- | --- |
|  | Main effects | p-value | Group-wise | ‘Healthy Aged’ | | ‘Aging Impaired’ | ‘AD Resilient’ | ‘AD Impaired |
| Linear Mixed Model + Tukey post hoc with LSD correction | Genotype | 0.949 | ‘Healthy Aged’ | x | | ***0.0379*** | 0.990 | 0.143 |
|  |  |  | ‘Aging Impaired’ | x | | x | 0.0671 | 0.975 |
|  | Cognitive Status | ***0.00192*** | ‘AD Resilient’ | x | | x | x | 0.226 |
|  |  |  | ‘AD Impaired | x | | x | x | x |
|  | Noise Floor: S1FL Inhibitory neurons | | | | | | | |
|  | Main effects | p-value | Group-wise | ‘Healthy Aged’ | | ‘Aging Impaired’ | ‘AD Resilient’ | ‘AD Impaired |
|  | Genotype | 0.643 | ‘Healthy Aged’ | x | | ***0.0238*** | 0.999 | 0.202 |
|  |  |  | ‘Aging Impaired’ | x | | x | ***0.0264*** | 0.848 |
|  | Cognitive Status | ***0.00137*** | ‘AD Resilient’ | x | | x | x | 0.229 |
|  |  |  | ‘AD Impaired | x | | x | x | x |
|  | Noise Floor: HPC | | | | | | | |
| Model | Main effects | p-values | | | | | | |
|  |  | Excitatory neurons | | | Inhibitory neurons | | | |
| Linear Mixed Model | Genotype | 0.638 | | | 0.642 | | | |
|  | Cognitive Status | 0.819 | | | 0.724 | | | |

**Supplementary Table 7.** Statistical analysis of SU channel noise floor; significance *p*<0.05; ‘Healthy Aged’: n=10; ‘Aging Impaired’: n=15; ‘AD Resilient’ rats: n=15; ‘AD Impaired’ rats: n =10. Significant p-values are bolded and italicized.

| Model | # of Activated SU: S1FL Excitatory neurons | | | | | | | |
| --- | --- | --- | --- | --- | --- | --- | --- | --- |
|  | Main effects | p-value | Group-wise | ‘Healthy Aged’ | | ‘Aging Impaired’ | ‘AD Resilient’ | ‘AD Impaired |
| Linear Mixed Model + Tukey post hoc with LSD correction | Genotype | 0.405 | ‘Healthy Aged’ | x | | 0.193 | 0.998 | 0.815 |
|  |  |  | ‘Aging Impaired’ | x | | x | 0.129 | 0.709 |
|  | Cognitive Status | ***0.0377*** | ‘AD Resilient’ | x | | x | x | 0.722 |
|  |  |  | ‘AD Impaired | x | | x | x | x |
| Model | # of Activated SU | | | | | | | |
|  | Main effects | p-values | | | | | | |
|  |  | S1FL Inhibitory neurons | | | HPC Excitatory neurons | | | |
| Linear Mixed Model | Genotype | 0.0667 | | | 0.303 | | | |
|  | Cognitive Status | 0.732 | | | 0.988 | | | |
| Model | # of Activated SU: HPC Inhibitory neurons | | | | | | | |
|  | Main effects | p-value | Group-wise | ‘Healthy Aged’ | | ‘Aging Impaired’ | ‘AD Resilient’ | ‘AD Impaired |
| Linear Mixed Model + Tukey post hoc with LSD correction | Genotype | ***0.0033*** | ‘Healthy Aged’ | x | | ***0.0168*** | 0.615 | 0.0766 |
|  |  |  | ‘Aging Impaired’ | x | | x | 0.210 | ***6.71e-5*** |
|  | Cognitive Status | 0.741 | ‘AD Resilient’ | x | | x | x | ***0.00624*** |
|  |  |  | ‘AD Impaired | x | | x | x | x |

**Supplementary Table 9.** Statistical analysis of the count of activated SU; significance *p*<0.05; 10 stimulation trials repetition in each animal subject; ‘Healthy Aged’: n=10; ‘Aging Impaired’: n=15; ‘AD Resilient’ rats: n=15; ‘AD Impaired’ rats: n =10. Significant p-values are bolded and italicized.

| Model | Evoked firing rate: S1FL Excitatory neurons | | | | | | | | | | |
| --- | --- | --- | --- | --- | --- | --- | --- | --- | --- | --- | --- |
|  | Main effects | p-value | Group-wise | | ‘Healthy Aged’ | | ‘Aging Impaired’ | | ‘AD Resilient’ | | ‘AD Impaired |
| Linear Mixed Model + Tukey post hoc with LSD correction | Genotype | 0.0795 | ‘Healthy Aged’ | | x | | ***0.00932*** | | 0.168 | | ***8.88e-4*** |
|  |  |  | ‘Aging Impaired’ | | x | | x | | 0.657 | | 0.543 |
|  | Cognitive Status | ***3.34e-4*** | ‘AD Resilient’ | | x | | x | | x | | 0.120 |
|  |  |  | ‘AD Impaired | | x | | x | | x | | x |
|  | Evoked firing rate: HPC Excitatory neurons | | | | | | | | | | |
|  | Main effects | p-value | Group-wise | | ‘Healthy Aged’ | | ‘Aging Impaired’ | | ‘AD Resilient’ | | ‘AD Impaired |
|  | Genotype | 0.103 | ‘Healthy Aged’ | | x | | 0.656 | | 0.219 | | 0.938 |
|  |  |  | ‘Aging Impaired’ | | x | | x | | ***0.0181*** | | 0.964 |
|  | Cognitive Status | ***0.0163*** | ‘AD Resilient’ | | x | | x | | x | | 0.104 |
|  |  |  | ‘AD Impaired | | x | | x | | x | | x |
|  | Evoked firing rate: HPC Inhibitory neurons | | | | | | | | | | |
|  | Main effects | p-value | Group-wise | | ‘Healthy Aged’ | | ‘Aging Impaired’ | | ‘AD Resilient’ | | ‘AD Impaired |
|  | Genotype | 0.468 | ‘Healthy Aged’ | | x | | 0.990 | | 0.985 | | ***0.0394*** |
|  |  |  | ‘Aging Impaired’ | | x | | x | | 0.919 | | ***0.0243*** |
|  | Cognitive Status | 0.118 | ‘AD Resilient’ | | x | | x | | x | | ***0.0378*** |
|  |  |  | ‘AD Impaired | | x | | x | | x | | x |
| Model | SU Firing Rates (Hz) | | | | | | | | | | |
|  | Main effects | p-values | | | | | | | | | |
|  |  | Baseline:  S1FL Excitatory neurons | | Baseline:  S1FL Inhibitory neurons | | Evoked:  S1FL Inhibitory neurons | | Baseline:  HPC Excitatory neurons | | Baseline:  HPC Inhibitory neurons | |
| Linear Mixed Model | Genotype | 0.187 | | 0.179 | | 0.224 | | 0.385 | | 0.977 | |
|  | Cognitive Status | 0.107 | | 0.191 | | 0.153 | | 0.413 | | 0.651 | |

**Supplementary Table 10.** Statistical analysis of the SU Firing rate; significance *p*<0.05; 10 stimulation trials repetition in each animal subject; ‘Healthy Aged’: n=10; ‘Aging Impaired’: n=15; ‘AD Resilient’ rats: n=15; ‘AD Impaired’ rats: n =10. Significant p-values are bolded and italicized.

| Model | 2D Δ Inter-Spike Interval (ΔISI) histograms: S1FL Excitatory Neurons | | | | |
| --- | --- | --- | --- | --- | --- |
|  | Group-wise | ‘Healthy Aged’ | ‘Aging Impaired’ | ‘AD Resilient’ | ‘AD Impaired |
| Permutation Test | ‘Healthy Aged’ | x | 0.482 | 0.716 | 0.839 |
|  | ‘Aging Impaired’ | x | x | 0.0600 | 0.2630 |
|  | ‘AD Resilient’ | x | x | x | 0.183 |
|  | ‘AD Impaired | x | x | x | x |
|  | 2D ΔISI histograms: S1FL Inhibitory Neurons | | | | |
|  | Group-wise | ‘Healthy Aged’ | ‘Aging Impaired’ | ‘AD Resilient’ | ‘AD Impaired |
|  | ‘Healthy Aged’ | x | 0.0810 | 0.379 | 0.243 |
|  | ‘Aging Impaired’ | x | x | 0.590 | 0.188 |
|  | ‘AD Resilient’ | x | x | x | 0.260 |
|  | ‘AD Impaired | x | x | x | x |
|  | 2D ΔISI histograms: HPC Excitatory Neurons | | | | |
|  | Group-wise | ‘Healthy Aged’ | ‘Aging Impaired’ | ‘AD Resilient’ | ‘AD Impaired |
|  | ‘Healthy Aged’ | x | 0.151 | 0.350 | 0.153 |
|  | ‘Aging Impaired’ | x | x | 0.127 | 0.217 |
|  | ‘AD Resilient’ | x | x | x | 0.115 |
|  | ‘AD Impaired | x | x | x | x |
|  | 2D ΔISI histograms: HPC Inhibitory Neurons | | | | |
|  | Group-wise | ‘Healthy Aged’ | ‘Aging Impaired’ | ‘AD Resilient’ | ‘AD Impaired |
|  | ‘Healthy Aged’ | x | 0.139 | 0.0790 | 0.423 |
|  | ‘Aging Impaired’ | x | x | 0.123 | 0.375 |
|  | ‘AD Resilient’ | x | x | x | 0.203 |
|  | ‘AD Impaired | x | x | x | x |

**Supplementary Table 11.** Statistical analysis of the mean 2D histograms of ISI; significance *p*<0.05; ‘Healthy Aged’: n=10; ‘Aging Impaired’: n=15; ‘AD Resilient’ rats: n=15; ‘AD Impaired’ rats: n =10. Significant p-values are bolded and italicized.

| Model | Irregular Firing: S1FL Excitatory neurons | | | | | | | | | | |
| --- | --- | --- | --- | --- | --- | --- | --- | --- | --- | --- | --- |
|  | Main effects | p-value | Group-wise | | ‘Healthy Aged’ | | ‘Aging Impaired’ | | ‘AD Resilient’ | | ‘AD Impaired |
| Linear Mixed Model + Tukey post hoc with LSD correction | Genotype | 0.803 | ‘Healthy Aged’ | | x | | 0.436 | | ***0.0328*** | | 0.408 |
|  |  |  | ‘Aging Impaired’ | | x | | x | | ***0.00268*** | | 0.814 |
|  | Cognitive Status | ***0.00689*** | ‘AD Resilient’ | | x | | x | | x | | ***0.0103*** |
|  |  |  | ‘AD Impaired | | x | | x | | x | | x |
|  | Irregular Firing: S1FL Inhibitory neurons | | | | | | | | | | |
|  | Main effects | p-value | Group-wise | | ‘Healthy Aged’ | | ‘Aging Impaired’ | | ‘AD Resilient’ | | ‘AD Impaired |
|  | Genotype | ***0.0282*** | ‘Healthy Aged’ | | x | | 0.322 | | 0.488 | | 0.257 |
|  |  |  | ‘Aging Impaired’ | | x | | x | | 0.0803 | | 0.370 |
|  | Cognitive Status | 0.583 | ‘AD Resilient’ | | x | | x | | x | | 0.603 |
|  |  |  | ‘AD Impaired | | x | | x | | x | | x |
|  | Irregular Firing: HPC Inhibitory neurons | | | | | | | | | | |
|  | Main effects | p-value | Group-wise | | ‘Healthy Aged’ | | ‘Aging Impaired’ | | ‘AD Resilient’ | | ‘AD Impaired |
|  | Genotype | 0.817 | ‘Healthy Aged’ | | x | | ***0.0175*** | | 0.901 | | 0.102 |
|  |  |  | ‘Aging Impaired’ | | x | | x | | ***0.0221*** | | 0.677 |
|  | Cognitive Status | ***0.00720*** | ‘AD Resilient’ | | x | | x | | x | | 0.122 |
|  |  |  | ‘AD Impaired | | x | | x | | x | | x |
| Model | Spiking Temporal Patterns | | | | | | | | | | |
|  | Main effects | p-values | | | | | | | | | |
|  |  | Bursty Firing: S1FL Excitatory Neurons | | Bursty Firing: S1FL Inhibitory Neurons | | Irregular Firing: HPC Excitatory Neurons | | Bursty Firing: HPC Excitatory Neurons | | Bursty Firing: HPC Inhibitory Neurons | |
| Linear Mixed Model | Genotype | 0.934 | | 0.163 | | 0.276 | | 0.232 | | 0.589 | |
|  | Cognitive Status | 0.363 | | 0.789 | | 0.938 | | 0.112 | | 0.238 | |

**Supplementary Table 12.** Statistical analysis of the percentage of neurons with irregular or bursty firing patterns; significance *p*<0.05; ‘Healthy Aged’: n=10; ‘Aging Impaired’: n=15; ‘AD Resilient’ rats: n=15; ‘AD Impaired’ rats: n =10. Significant p-values are bolded and italicized.

| NEI: S1FL Excitatory neurons | | | | | | | | | | | | | | | | |
| --- | --- | --- | --- | --- | --- | --- | --- | --- | --- | --- | --- | --- | --- | --- | --- | --- |
| Model | | Main effects | p-value | | | Group-wise | | | ‘Healthy Aged’ | | | | ‘Aging Impaired’ | ‘AD Resilient’ | | ‘AD Impaired |
| Linear Mixed Model + Tukey post hoc with LSD correction | | Genotype | 0.572 | | | ‘Healthy Aged’ | | | x | | | | ***6.93e-7*** | 0.122 | | ***1.53e-4*** |
|  |  |  |  |  |  | ‘Aging Impaired’ | | | x | | | | x | ***0.00211*** | | 0.530 |
|  |  | Cognitive Status | ***2.19e-7*** | | | ‘AD Resilient’ | | | x | | | | x | x | | ***0.0302*** |
|  |  |  |  |  |  | ‘AD Impaired | | | x | | | | x | x | | x |
| NEI | | | | | | | | | | | | | | | | |
| Model | | Main effects | | p-values | | | | | | | | | | | | |
|  |  |  |  | S1FL: Inhibitory neurons | | | | | | | HPC: Excitatory neurons | | | | | |
| Linear Mixed Model | | Genotype | | 0.325 | | | | | | | 0.519 | | | | | |
|  |  | Cognitive Status | | 0.842 | | | | | | | 0.998 | | | | | |
| NEI: HPC Inhibitory neurons | | | | | | | | | | | | | | | | |
| Model | Main effects | | p-value | | Group-wise | | | ‘Healthy Aged’ | | | | ‘Aging Impaired’ | | ‘AD Resilient’ | | ‘AD Impaired |
| Linear Mixed Model + Tukey post hoc with LSD correction | Genotype | | 0.255 | | ‘Healthy Aged’ | | | x | | | | 0.531 | | 0.204 | | 0.866 |
|  |  |  |  |  | ‘Aging Impaired’ | | | x | | | | x | | ***0.00982*** | | 0.437 |
|  | Cognitive Status | | ***0.0386*** | | ‘AD Resilient’ | | | x | | | | x | | x | | ***0.0272*** |
|  |  |  |  |  | ‘AD Impaired | | | x | | | | x | | x | | x |
| Model | | # of Activated SU per Stimulus Pulse | | | | | | | | | | | | | | |
|  |  | Main effects | p-values | | | | | | | | | | | | | |
|  |  |  | S1FL Excitatory Neurons | | | | S1FL Inhibitory Neurons | | | HPC Excitatory Neurons | | | | | HPC Inhibitory Neurons | |
| Linear Mixed Model | | Genotype | 0.232 | | | | 0.379 | | | 0.935 | | | | | 0.0952 | |
|  |  | Cognitive Status | 0.199 | | | | 0.748 | | | 0.976 | | | | | 0.655 | |
|  |  | # of Total Activated SU per Stimulation Trial | | | | | | | | | | | | | | |
|  |  | Main effects | p-values | | | | | | | | | | | | | |
|  |  |  | S1FL Excitatory Neurons | | | | S1FL Inhibitory Neurons | | | HPC Excitatory Neurons | | | | | HPC Inhibitory Neurons | |
|  |  | Genotype | 0.123 | | | | 0.0655 | | | 0.699 | | | | | 0.0786 | |
|  |  | Cognitive Status | 0.191 | | | | 0.685 | | | 0.754 | | | | | 0.523 | |

**Supplementary Table 13.** Statistical analysis of Neuronal Engagement Index (NEI), # of activated SU per stimulus pulse, and # of total activated SU per stimulation trial; significance *p*<0.05; ‘Healthy Aged’: n=10; ‘Aging Impaired’: n=15; ‘AD Resilient’ rats: n=15; ‘AD Impaired’ rats: n =10. Significant p-values are bolded and italicized.

| Model | # of CCG-based synaptic connection pairs: S1FL | | | | | | |
| --- | --- | --- | --- | --- | --- | --- | --- |
|  | Main effects | p-values | | | | | |
| Linear Mixed Model | Genotype | 0.486 | | | | | |
|  | Cognitive Status | 0.891 | | | | | |
| Model | # of CCG-based synaptic connection pairs: HPC | | | | | | |
|  | Main effects | p-value | Group-wise | ‘Healthy Aged’ | ‘Aging Impaired’ | ‘AD Resilient’ | ‘AD Impaired |
| Linear Mixed Model + Tukey post hoc with LSD correction | Genotype | 0.490 | ‘Healthy Aged’ | x | 0.218 | 0.958 | 0.205 |
|  |  |  | ‘Aging Impaired’ | x | x | 0.476 | ***0.00418*** |
|  | Cognitive Status | 0.313 | ‘AD Resilient’ | x | x | x | ***0.0130*** |
|  |  |  | ‘AD Impaired | x | x | x | x |

**Supplementary Table 14.** Statistical analysis of the total count of CCG-based connection pairs; significance *p*<0.05; ‘Healthy Aged’: n=10; ‘Aging Impaired’: n=15; ‘AD Resilient’ rats: n=15; ‘AD Impaired’ rats: n =10. Significant p-values are bolded and italicized.

| Model | Percentage of Excitatory CCG-based connections: S1FL | | | | | | |
| --- | --- | --- | --- | --- | --- | --- | --- |
|  | Main effects | p-value | Group-wise | ‘Healthy Aged’ | ‘Aging Impaired’ | ‘AD Resilient’ | ‘AD Impaired |
| Linear Mixed Model + Tukey post hoc with LSD correction | Genotype | 0.358 | ‘Healthy Aged’ | x | 0.217 | 0.698 | 0.127 |
|  |  |  | ‘Aging Impaired’ | x | x | 0.376 | 0.400 |
|  | Cognitive Status | ***0.0419*** | ‘AD Resilient’ | x | x | x | 0.0728 |
|  |  |  | ‘AD Impaired | x | x | x | x |
|  | Percentage of Excitatory CCG-based connections: HPC | | | | | | |
|  | Main effects | p-value | Group-wise | ‘Healthy Aged’ | ‘Aging Impaired’ | ‘AD Resilient’ | ‘AD Impaired |
|  | Genotype | 0.583 | ‘Healthy Aged’ | x | ***0.0152*** | 0.0735 | 0.176 |
|  |  |  | ‘Aging Impaired’ | x | x | 0.538 | 0.289 |
|  | Cognitive Status | 0.143 | ‘AD Resilient’ | x | x | x | 0.660 |
|  |  |  | ‘AD Impaired | x | x | x | x |

**Supplementary Table 15.** Statistical analysis of the percentage of excitatory CCG-based connections; significance *p*<0.05; ‘Healthy Aged’: n=10; ‘Aging Impaired’: n=15; ‘AD Resilient’ rats: n=15; ‘AD Impaired’ rats: n =10. Significant p-values are bolded and italicized.

| Model | Percentage of E-E vs. E-I connections: S1FL | | | | | | |
| --- | --- | --- | --- | --- | --- | --- | --- |
|  | Main effects | p-values | | | | | |
| Linear Mixed Model | Genotype | 0.467 | | | | | |
|  | Cognitive Status | 0.0665 | | | | | |
| Model | Percentage of E-E vs. E-I connections: HPC | | | | | | |
|  | Main effects | p-value | Group-wise | ‘Healthy Aged’ | ‘Aging Impaired’ | ‘AD Resilient’ | ‘AD Impaired |
| Linear Mixed Model + Tukey post hoc with LSD correction | Genotype | ***0.00400*** | ‘Healthy Aged’ | x | 0.102 | 0.0942 | 0.401 |
|  |  |  | ‘Aging Impaired’ | x | x | ***0.00337*** | ***0.0226*** |
|  | Cognitive Status | 0.0677 | ‘AD Resilient’ | x | x | x | 0.393 |
|  |  |  | ‘AD Impaired | x | x | x | x |

**Supplementary Table 16.** Statistical analysis of the percentage of E-E vs. E-I connections over the total excitatory CCG-based connections; significance *p*<0.05; ‘Healthy Aged’: n=10; ‘Aging Impaired’: n=15; ‘AD Resilient’ rats: n=15; ‘AD Impaired’ rats: n =10. Significant p-values are bolded and italicized.

| Model | 2Strength of I connections: S1FL | | | | | | | |
| --- | --- | --- | --- | --- | --- | --- | --- | --- |
|  | Main effects | p-value | Group-wise | ‘Healthy Aged’ | | ‘Aging Impaired’ | ‘AD Resilient’ | ‘AD Impaired |
| Linear Mixed Model + Tukey post hoc with LSD correction | Genotype | 0.361 | ‘Healthy Aged’ | x | | ***0.00245*** | 0.465 | ***0.00265*** |
|  |  |  | ‘Aging Impaired’ | x | | x | ***0.0125*** | 0.615 |
|  | Cognitive Status | ***9.53e-05*** | ‘AD Resilient’ | x | | x | x | ***0.0111*** |
|  |  |  | ‘AD Impaired | x | | x | x | x |
|  | Strength of I connections: HPC | | | | | | | |
|  | Main effects | p-value | Group-wise | ‘Healthy Aged’ | | ‘Aging Impaired’ | ‘AD Resilient’ | ‘AD Impaired |
|  | Genotype | 0.427 | ‘Healthy Aged’ | x | | 0.467 | 0.155 | 0.0777 |
|  |  |  | ‘Aging Impaired’ | x | | x | 0.510 | ***0.0251*** |
|  | Cognitive Status | 0.0741 | ‘AD Resilient’ | x | | x | x | ***0.00523*** |
|  |  |  | ‘AD Impaired | x | | x | x | x |
| Model | Strength of E connections | | | | | | | |
|  | Main effects | p-values | | | | | | |
|  |  | S1FL | | | HPC | | | |
| Linear Mixed Model | Genotype | 0.646 | | | 0.251 | | | |
|  | Cognitive Status | 0.211 | | | 0.425 | | | |

**Supplementary Table 17.** Statistical analysis of the CCG-based connection strength; significance *p*<0.05; ‘Healthy Aged’: n=10; ‘Aging Impaired’: n=15; ‘AD Resilient’ rats: n=15; ‘AD Impaired’ rats: n =10. Significant p-values are bolded and italicized.

| Model | Percentage of I-E vs. I-I connections: S1FL | | | | | | | |
| --- | --- | --- | --- | --- | --- | --- | --- | --- |
|  | Main effects | p-value | | Group-wise | ‘Healthy Aged’ | ‘Aging Impaired’ | ‘AD Resilient’ | ‘AD Impaired |
| Linear Mixed Model + Tukey post hoc with LSD correction | Genotype | 0.265 | | ‘Healthy Aged’ | x | ***0.0367*** | 0.959 | 0.720 |
|  |  |  |  | ‘Aging Impaired’ | x | x | ***0.0312*** | 0.140 |
|  | Cognitive Status | ***0.00907*** | | ‘AD Resilient’ | x | x | x | 0.744 |
|  |  |  |  | ‘AD Impaired | x | x | x | x |
| Model | Percentage of I-E vs. I-I connections: HPC | | | | | | | |
|  | Main effects | | p-values | | | | | |
| Linear Mixed Model | Genotype | | 0.643 | | | | | |
|  | Cognitive Status | | 0.536 | | | | | |

**Supplementary Table 16.** Statistical analysis of the percentage of I-E vs. I-I connections over the total inhibitory CCG-based connections; significance *p*<0.05; ‘Healthy Aged’: n=10; ‘Aging Impaired’: n=15; ‘AD Resilient’ rats: n=15; ‘AD Impaired’ rats: n =10. Significant p-values are bolded and italicized.

| Model | Count of Postsynaptic E neurons by Excitatory Convergence Degree: S1FL Bin >5 | | | | | | | |
| --- | --- | --- | --- | --- | --- | --- | --- | --- |
|  | Main effects | p-value | Group-wise | ‘Healthy Aged’ | | ‘Aging Impaired’ | ‘AD Resilient’ | ‘AD Impaired |
| Linear Mixed Model + Tukey post hoc with LSD correction | Genotype | 0.749 | ‘Healthy Aged’ | x | | 0.116 | 0.962 | 0.397 |
|  |  |  | ‘Aging Impaired’ | x | | x | 0.421 | 0.828 |
|  | Cognitive Status | ***0.0302*** | ‘AD Resilient’ | x | | x | x | 0.197 |
|  |  |  | ‘AD Impaired | x | | x | x | x |
| Model | Count of Postsynaptic E neurons by Excitatory Convergence Degree | | | | | | | |
|  | Main effects | p-values | | | | | | |
|  |  | S1FL: Bin =1 | | | S1FL: Bin 2-5 | | | |
| Linear Mixed Model | Genotype | 0.774 | | | 0.816 | | | |
|  | Cognitive Status | 0.141 | | | 0.202 | | | |
| Model | Count of Postsynaptic E neurons by Excitatory Convergence Degree: HPC Bin 2-5 | | | | | | | |
|  | Main effects | p-value | Group-wise | ‘Healthy Aged’ | | ‘Aging Impaired’ | ‘AD Resilient’ | ‘AD Impaired |
| Linear Mixed Model + Tukey post hoc with LSD correction | Genotype | 0.219 | ‘Healthy Aged’ | x | | 0.591 | 0.726 | 0.913 |
|  |  |  | ‘Aging Impaired’ | x | | x | 0.149 | 0.922 |
|  | Cognitive Status | ***0.0352*** | ‘AD Resilient’ | x | | x | x | 0.369 |
|  |  |  | ‘AD Impaired | x | | x | x | x |
| Model | Count of Postsynaptic E neurons by Excitatory Convergence Degree | | | | | | | |
|  | Main effects | p-values | | | | | | |
|  |  | S1FL: Bin =1 | | | S1FL: Bin >5 | | | |
| Linear Mixed Model | Genotype | 0.219 | | | 0.791 | | | |
|  | Cognitive Status | 0.0353 | | | 0.341 | | | |

**Supplementary Table 17.** Statistical analysis of the count of postsynaptic E neurons by excitatory convergence degree; significance *p*<0.05; ‘Healthy Aged’: n=10; ‘Aging Impaired’: n=15; ‘AD Resilient’ rats: n=15; ‘AD Impaired’ rats: n =10. Significant p-values are bolded and italicized.

| Model | Count of Postsynaptic I neurons by Excitatory Convergence Degree | | | | | | | | |
| --- | --- | --- | --- | --- | --- | --- | --- | --- | --- |
|  | Main effects | p-values | | | | | | | |
|  |  | S1FL: Bin =1 | | S1FL: Bin 2-5 | | | S1FL: Bin >5 | | |
| Linear Mixed Model | Genotype | 0.332 | | 0.160 | | | 0.970 | | |
|  | Cognitive Status | 0.539 | | 0.456 | | | 0.114 | | |
| Model | Count of Postsynaptic I neurons by Excitatory Convergence Degree: HPC Bin 2-5 | | | | | | | | |
|  | Main effects | p-value | Group-wise | | ‘Healthy Aged’ | ‘Aging Impaired’ | | ‘AD Resilient’ | ‘AD Impaired |
| Linear Mixed Model + Tukey post hoc with LSD correction | Genotype | ***0.00597*** | ‘Healthy Aged’ | | x | 0.899 | | 0.573 | ***0.0107*** |
|  |  |  | ‘Aging Impaired’ | | x | x | | 0.949 | 0.0724 |
|  | Cognitive Status | ***0.0387*** | ‘AD Resilient’ | | x | x | | x | 0.168 |
|  |  |  | ‘AD Impaired | | x | x | | x | x |
|  | Count of Postsynaptic I neurons by Excitatory Convergence Degree: HPC Bin >5 | | | | | | | | |
|  | Main effects | p-value | Group-wise | | ‘Healthy Aged’ | ‘Aging Impaired’ | | ‘AD Resilient’ | ‘AD Impaired |
|  | Genotype | 0.182 | ‘Healthy Aged’ | | x | 0.625 | | 0.500 | 0.875 |
|  |  |  | ‘Aging Impaired’ | | x | x | | 0.0868 | 0.972 |
|  | Cognitive Status | ***0.0196*** | ‘AD Resilient’ | | x | x | | x | 0.199 |
|  |  |  | ‘AD Impaired | | x | x | | x | x |
| Model | Count of Postsynaptic I neurons by Excitatory Convergence Degree: HPC Bin =1 | | | | | | | | |
|  | Main effects | p-values | | | | | | | |
| Linear Mixed Model | Genotype | 0.377 | | | | | | | |
|  | Cognitive Status | 0.0741 | | | | | | | |

**Supplementary Table 18.** Statistical analysis of the count of postsynaptic I neurons by excitatory convergence degree; significance *p*<0.05; ‘Healthy Aged’: n=10; ‘Aging Impaired’: n=15; ‘AD Resilient’ rats: n=15; ‘AD Impaired’ rats: n =10. Significant p-values are bolded and italicized.

| Model | Count of Presynaptic I neurons by Divergence Degree onto E Neurons: S1FL Bin 2-5 | | | | | | | |
| --- | --- | --- | --- | --- | --- | --- | --- | --- |
|  | Main effects | p-value | Group-wise | ‘Healthy Aged’ | | ‘Aging Impaired’ | ‘AD Resilient’ | ‘AD Impaired |
| Linear Mixed Model + Tukey post hoc with LSD correction | Genotype | 0.238 | ‘Healthy Aged’ | x | | ***0.0457*** | 0.0818 | ***0.0371*** |
|  |  |  | ‘Aging Impaired’ | x | | x | 0.940 | 0.779 |
|  | Cognitive Status | 0.202 | ‘AD Resilient’ | x | | x | x | 0.741 |
|  |  |  | ‘AD Impaired | x | | x | x | x |
| Model | Count of Presynaptic I neurons by Divergence Degree onto E Neurons | | | | | | | |
|  | Main effects | p-values | | | | | | |
|  |  | S1FL: Bin =1 | | | S1FL: Bin >5 | | | |
| Linear Mixed Model | Genotype | 0.601 | | | 0.589 | | | |
|  | Cognitive Status | 0.103 | | | 0.381 | | | |
|  | Count of Presynaptic I neurons by Divergence Degree onto E Neurons | | | | | | | |
|  | Main effects | p-values | | | | | | |
|  |  | HPC: Bin =1 | | | HPC: Bin 2-5 | | | |
|  | Genotype | 0.648 | | | 0.0912 | | | |
|  | Cognitive Status | 0.901 | | | 0.278 | | | |

**Supplementary Table 19.** Statistical analysis of the count of presynaptic I neurons by divergence degree onto E neurons; significance *p*<0.05; ‘Healthy Aged’: n=10; ‘Aging Impaired’: n=15; ‘AD Resilient’ rats: n=15; ‘AD Impaired’ rats: n =10. Significant p-values are bolded and italicized.

| Model | Count of Presynaptic I neurons by Divergence Degree onto I Neurons: S1FL Bin 2-5 | | | | | | | |
| --- | --- | --- | --- | --- | --- | --- | --- | --- |
|  | Main effects | p-value | Group-wise | ‘Healthy Aged’ | ‘Aging Impaired’ | | ‘AD Resilient’ | ‘AD Impaired |
| Linear Mixed Model + Tukey post hoc with LSD correction | Genotype | 0.146 | ‘Healthy Aged’ | x | 0.0833 | | 0.524 | 0.742 |
|  |  |  | ‘Aging Impaired’ | x | x | | ***0.0157*** | 0.180 |
|  | Cognitive Status | ***0.0417*** | ‘AD Resilient’ | x | x | | x | 0.341 |
|  |  |  | ‘AD Impaired | x | x | | x | x |
| Model | Count of Presynaptic I neurons by Divergence Degree onto I Neurons: S1FL Bin =1 | | | | | | | |
|  | Main effects | p-values | | | | | | |
| Linear Mixed Model | Genotype | 0.0693 | | | | | | |
|  | Cognitive Status | 0.388 | | | | | | |
|  | Count of Presynaptic I neurons by Divergence Degree onto I Neurons | | | | | | | |
|  | Main effects | p-values | | | | | | |
|  |  | HPC: Bin =1 | | | | HPC: Bin 2-5 | | |
|  | Genotype | 0.955 | | | | 0.455 | | |
|  | Cognitive Status | 0.892 | | | | 0.281 | | |

**Supplementary Table 19.** Statistical analysis of the count of presynaptic I neurons by divergence degree onto I neurons; significance *p*<0.05; ‘Healthy Aged’: n=10; ‘Aging Impaired’: n=15; ‘AD Resilient’ rats: n=15; ‘AD Impaired’ rats: n =10. Significant p-values are bolded and italicized.

| S1FL: Excitatory neurons | | | | | | Learning |
| --- | --- | --- | --- | --- | --- | --- |
| Model | Variable | | | | | p-value |
| ANCOVA | Degree of Excitatory Convergence | | | | | ***6.33e-17*** |
|  | Genotype | | | | | 0.650 |
|  | Cognitive Status | | | | | 0.164 |
|  | Degree of Excitatory Convergence: Genotype | | | | | 0.0502 |
|  | Degree of Excitatory Convergence: Cognitive Status | | | | | 0.355 |
|  | Genotype: Cognitive Status | | | | | 0.0630 |
|  | Degree of Excitatory Convergence: Genotype: Cognitive Status | | | | | ***0.000469*** |
| Pairwise Post hoc Slope Comparisons: S1FL Excitatory neurons | | | | | | |
| Group-wise | | ‘Healthy Aged’ | ‘Aging Impaired’ | ‘AD Resilient’ | ‘AD Impaired | |
| ‘Healthy Aged’ | | x | ***7.17e-5*** | ***0.00184*** | 0.712 | |
| ‘Aging Impaired’ | | x | x | 0.408 | ***0.00113*** | |
| ‘AD Resilient’ | | x | x | x | ***0.000471*** | |
| ‘AD Impaired | | x | x | x | x | |

| S1FL: Inhibitory neurons | | | | | | Learning |
| --- | --- | --- | --- | --- | --- | --- |
| Model | Variable | | | | | p-value |
| ANCOVA | Degree of Excitatory Convergence | | | | | ***1.94e-05*** |
|  | Genotype | | | | | 0.260 |
|  | Cognitive Status | | | | | 0.435 |
|  | Degree of Excitatory Convergence: Genotype | | | | | 0.209 |
|  | Degree of Excitatory Convergence: Cognitive Status | | | | | 0.157 |
|  | Genotype: Cognitive Status | | | | | ***0.000339*** |
|  | Degree of Excitatory Convergence: Genotype: Cognitive Status | | | | | ***0.00239*** |
| Pairwise Post hoc Slope Comparisons: S1FL Inhibitory neurons | | | | | | |
| Group-wise | | ‘Healthy Aged’ | ‘Aging Impaired’ | ‘AD Resilient’ | ‘AD Impaired | |
| ‘Healthy Aged’ | | x | ***0.00323*** | ***0.00231*** | 0.0860 | |
| ‘Aging Impaired’ | | x | x | 0.462 | ***0.0272*** | |
| ‘AD Resilient’ | | x | x | x | 0.125 | |
| ‘AD Impaired | | x | x | x | x | |

| HPC: Excitatory neurons | | | | | | Learning |
| --- | --- | --- | --- | --- | --- | --- |
| Model | Variable | | | | | p-value |
| ANCOVA | Degree of Excitatory Convergence | | | | | ***3.72e-14*** |
|  | Genotype | | | | | ***0.0343*** |
|  | Cognitive Status | | | | | 0.0867 |
|  | Degree of Excitatory Convergence: Genotype | | | | | 0.923 |
|  | Degree of Excitatory Convergence: Cognitive Status | | | | | 0.748 |
|  | Genotype: Cognitive Status | | | | | 0.348 |
|  | Degree of Excitatory Convergence: Genotype: Cognitive Status | | | | | 0.355 |
| Pairwise Post hoc Slope Comparisons: HPC Excitatory neurons | | | | | | |
| Group-wise | | ‘Healthy Aged’ | ‘Aging Impaired’ | ‘AD Resilient’ | ‘AD Impaired | |
| ‘Healthy Aged’ | | x | 0.399 | 0.429 | 0.638 | |
| ‘Aging Impaired’ | | x | x | 0.768 | 0.677 | |
| ‘AD Resilient’ | | x | x | x | 0.599 | |
| ‘AD Impaired | | x | x | x | x | |

| HPC: Inhibitory neurons | | | | | | Learning |
| --- | --- | --- | --- | --- | --- | --- |
| Model | Variable | | | | | p-value |
| ANCOVA | Degree of Excitatory Convergence | | | | | ***3.66e-13*** |
|  | Genotype | | | | | 0.183 |
|  | Cognitive Status | | | | | ***0.000372*** |
|  | Degree of Excitatory Convergence: Genotype | | | | | 0.885 |
|  | Degree of Excitatory Convergence: Cognitive Status | | | | | 0.953 |
|  | Genotype: Cognitive Status | | | | | 0.888 |
|  | Degree of Excitatory Convergence: Genotype: Cognitive Status | | | | | ***0.0235*** |
| Pairwise Post hoc Slope Comparisons: HPC Inhibitory neurons | | | | | | |
| Group-wise | | ‘Healthy Aged’ | ‘Aging Impaired’ | ‘AD Resilient’ | ‘AD Impaired | |
| ‘Healthy Aged’ | | x | ***0.0411*** | 0.0704 | 0.989 | |
| ‘Aging Impaired’ | | x | x | 0.174 | 0.201 | |
| ‘AD Resilient’ | | x | x | x | ***0.0258*** | |
| ‘AD Impaired | | x | x | x | x | |

**Supplementary Table 20.** Statistical analysis of the relationship between the degree of excitatory convergence and NEI; significance *p*<0.05; ‘Healthy Aged’: n=10; ‘Aging Impaired’: n=15; ‘AD Resilient’ rats: n=15; ‘AD Impaired’ rats: n =10. Significant p-values are bolded and italicized.
